# An NKX-COUP-TFII genomic code for mucosal vascular addressins and organ morphogenesis

**DOI:** 10.1101/2022.02.17.480956

**Authors:** Thanh Theresa Dinh, Menglan Xiang, Anusha Rajaraman, Yongzhi Wang, Nicole Salazar, Walter Roper, Siyeon Rhee, Kevin Brulois, Ed O’Hara, Helena Kiefel, Truc Dinh, Yuhan Bi, Dalila Gonzalez, Evan Bao, Kristy Red-Horse, Peter Balogh, Fanni Gábris, Balázs Gaszner, Gergely Berta, Junliang Pan, Eugene C. Butcher

**Affiliations:** Laboratory of Immunology and Vascular Biology, Department of Pathology, Stanford University School of Medicine, Stanford, CA, USA; Palo Alto Veterans Institute for Research, Palo Alto, CA, USA; Department of Molecular Cell Biology and Immunology, Vrije Universiteit Medical Center, Amsterdam, The Netherlands; Department of Clinical Science Malmo, Section of Surgery, Lund University, Malmo, Sweden; Columbia University Vagelos College of Physicians and Surgeons; Department of Biology, Stanford University, Stanford, CA, USA; University of California, San Diego, La Jolla, CA, USA; Department of Immunology and Biotechnology, University of Pécs Medical School, Pécs Hungary; Lymphoid Organogenesis Research Team, Szentágothai Research Center, Pécs, Hungary; Department of Anatomy, University of Pécs Medical School, Pécs, Hungary; Department of Medical Biology and Central Electron Microscopy Laboratory, University of Pécs Medical School, Pécs, Hungary; The Center for Molecular Biology and Medicine, Veterans Affairs Palo Alto Health Care System, Palo Alto, CA, USA

**Author notes:** These authors contributed equally: Thanh Theresa Dinh, Menglan Xiang and Anusha Rajaraman. Equal contribution (Junliang Pan, Eugene C. Butcher). Lead contact: Eugene C. Butcher.

## Abstract

Immunoglobulin family and carbohydrate vascular addressins encoded by *Madcam1* and *St6gal1* control lymphocyte homing into intestinal tissues, regulating immunity and inflammation. The addressins are developmentally programmed to decorate endothelial cells lining gut post-capillary and high endothelial venules, providing a prototypical example of organ- and segment-specific endothelial specialization. We identify conserved NKX-COUP-TFII composite elements (NCCE) in regulatory regions of *Madcam1* and *St6gal1* that bind intestinal homeodomain protein NKX2-3 cooperatively with venous nuclear receptor COUP-TFII to activate transcription. The *Madcam1* element also integrates repressive signals from arterial/capillary Notch effectors. Pan-endothelial COUP-TFII overexpression induces ectopic addressin expression in NKX2-3^+^ capillaries, while NKX2-3 deficiency abrogates expression by HEV. Phylogenetically conserved NCCE are enriched in genes involved in neuron migration and morphogenesis of the heart, kidney, pancreas and other organs. Our results define a genomic address code for targeted expression of mucosal vascular addressins and implicate NCCE in fundamental processes in cell specification and development.

## INTRODUCTION

The blood vasculature is segmentally specialized for physiological functions specific to arterial, capillary, venous and sinus vessels, and regionally specialized for functions specific to different organs, tissues or microenvironments (Aird, 2007a; b). A prototypical example is the segmental and organotypic specialization of vascular endothelium for control of lymphocyte and immune cell traffic (Berg et al., 1989). Blood borne leukocytes adhere and extravasate preferentially through vascular segments immediately downstream of capillaries: postcapillary venules (PCV) in sites of inflammation and immune surveillance, and specialized high endothelial venules (HEV) in secondary and tertiary lymphoid tissues (Ager, 2017; Girard et al., 2012; Hayasaka et al., 2010; von Andrian and Mempel, 2003). HEV and PCV in sites of immune surveillance display organ-specific adhesion molecules termed vascular addressins that direct tissue-specific lymphocyte homing in support of regional immune specialization and responses (Ager, 2017; Butcher and Picker, 1996) (Berg *et al*., 1989). The mucosal vascular addressin-1, MAdCAM1, is an Ig and mucin family member expressed by HEV in gut associated lymphoid tissues (GALT) and by venules for lymphocyte homing into the intestinal lamina propria; but not normally by HEV in other tissues in the adult (Streeter et al., 1988) (Briskin et al., 1993) (Briskin et al., 1997). MAdCAM1 recruits T and B cells bearing the gut homing receptor integrin α_4_β_7_. Intestinal HEV and PCV are also decorated by α2-6 sialic acid-capped glycotopes recognized by the B cell specific lectin CD22 (Siglec2) (Lee et al., 2014b). These glycotopes constitute a B cell-specific mucosal vascular addressin, designated here BMAd, which selectively enhances the homing of B cells into GALT (Lee et al., 2014b) where they contribute to the IgA response to intestinal antigens and pathogens (Ballet et al., 2021). BMAd is synthesized by beta-galactoside alpha-2,6-sialyltransferase 1, encoded by *St6gal1*, which like *Madcam1* is selectively expressed by Peyer’s patch HEV in the GI tract (Lee *et al*., 2014b). While cis-acting composite motifs that confer endothelial-specific gene expression have been defined (De Val et al., 2008), the genomic motifs and transcriptional control mechanisms that direct the venule- and intestine-specific expression of the addressins remain to be elucidated. The intestinal homeodomain protein NKX2-3 is required for normal expression of MAdCAM1 by PCV and HEV in the adult (Czömpöly et al., 2011; Pabst et al., 2000; Wang et al., 2000), but how this pan-intestinal factor is recruited for selective venous expression is unclear, and the transcriptional control of *St6gal1* in EC has not been studied.

Here we describe a short genomic element that helps direct both the organ- and segment-specific expression of these mucosal vascular addressins. We identify composite elements (CE) in conserved regulatory regions of *Madcam1* and *St6gal1* that combine binding sites for NKX2-3 and for the master venous transcription factor (TF) COUP-TFII, an orphan nuclear hormone receptor expressed by venous but not capillary or arterial EC. We show that these NKX-COUP-TFII composite elements (NCCE) seed cooperative binding and heterodimerization of NKX2-3 with COUP-TFII, creating a transcriptional activation complex that promotes addressin expression. We also identify an E-box in the *Madcam1* CE that binds to the capillary/arterial transcriptional repressor HEY1 to suppress *Madcam1* expression. Phylogenetically conserved NCCE genomic motifs are enriched in association with genes involved in embryonic organ morphogenesis, including genes implicated in COUP-TFII- and NKX2-5, NKX2-2 or NKX2-1-dependent cardiovascular, pancreatic and neuronal specification. Thus our results show that COUP-TFII forms transcriptional activation complexes with NKX homeodomain partners on NCCE to target the vascular addressins, and implicate NCCE in embryonic organogenesis and cell specification.

## RESULTS

### A conserved composite element (CE) in the *Madcam1* promoter binds NKX2-3, COUP-TFII and HEY1

To identify regulatory sequences governing MAdCAM1 targeting to gut associated HEV, we initially applied Assay for Transposase-Accessible Chromatin using sequencing (ATAC-seq) to sorted blood high (HEC) and capillary endothelial cells (CapEC) from Peyer’s patches (PP) and peripheral lymph nodes (PLN). ATAC-seq identified a region near the *Madcam1* transcription start site that is transposon-accessible in PP HECs but not in MAdCAM1-negative PLN HEC or PP CapEC (Figure 1A). This region encompasses a previously described proximal NF-κB binding site (Sampaio et al., 1995; Wagner et al., 1996), and a nearby combinatorial motif consisting of a phylogenetically conserved candidate homeodomain (HD) and nuclear receptor binding sites, potential binding sites for gut HEV-expressed NKX2-3 and COUP-TFII (encoded by *Nr2f2*), respectively (Figure 1B and S1). Two canonical COUP-TFII ‘half sites’ are spaced seven nucleotides apart in the mouse (TGACCC, highlighted in green in Figure 1B and 1C), a spacing consistent with COUP-TFII homodimer binding (Cooney et al., 1992), with the COUP-TFII motif within the “A” site overlapping the homeodomain motif (purple line and text, Figure 1B, 1C, and S1). Interestingly, a canonical E-box motif (red line) overlaps both the homeodomain and the “A” site COUP-TFII motif (Figure 1B and C). E-box motifs bind Notch downstream transcriptional repressors HEY1 and HES1 (Fischer and Gessler, 2007), which are constitutively expressed in CapEC and arterial EC ((Lee et al., 2014a) and Figure S1). Overlap of the E-box and COUP-TFII binding motifs suggested the potential for competition between capillary and venular TFs. We refer to the overall sequence hereafter as the *Madcam1* composite element (CE).

**Figure 1.**
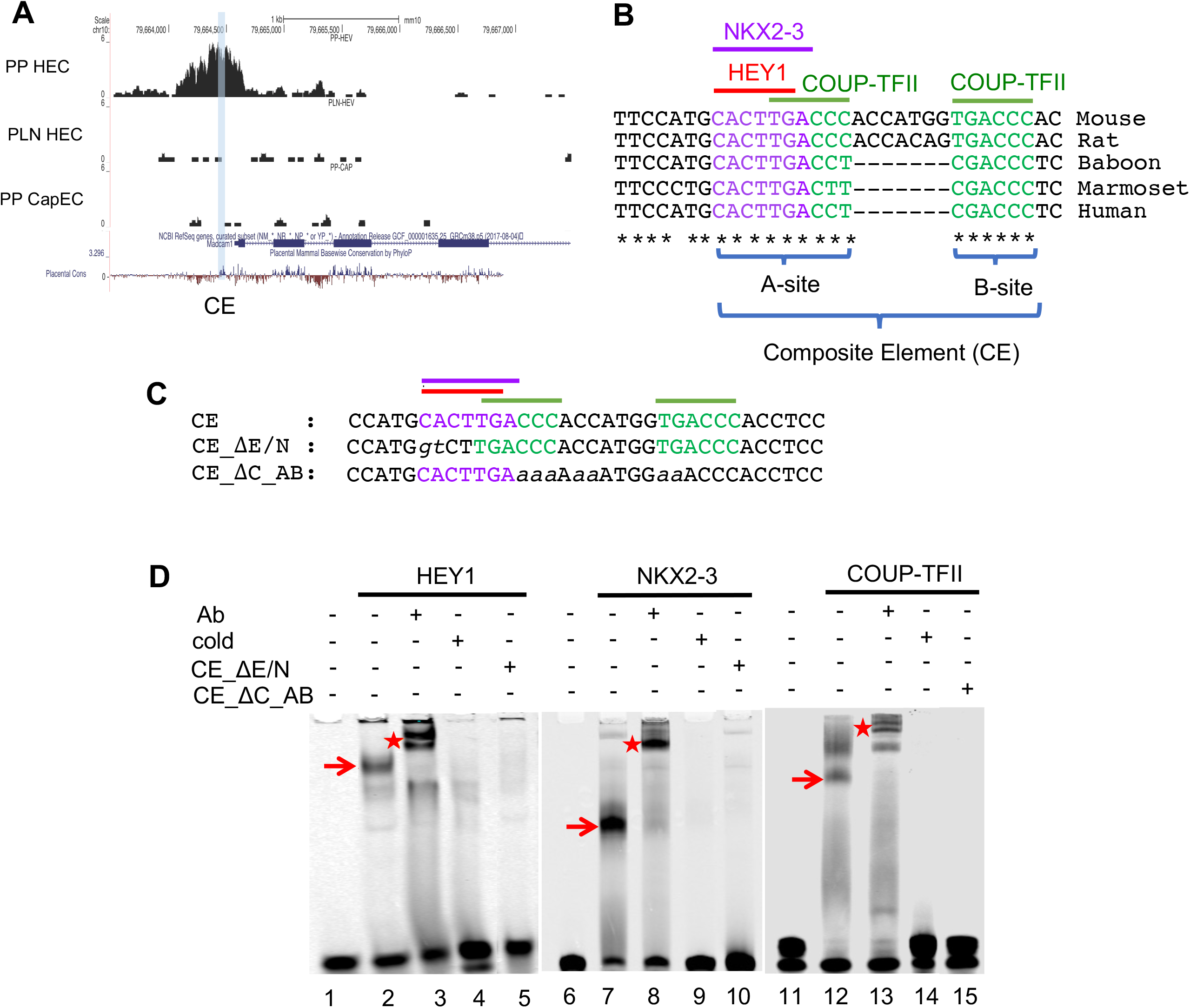
An evolutionarily conserved composite element (CE) in the *Madcam1* promoter binds NKX2-3, COUP-TFII and HEY1. (A) Normalized ATAC-seq signal in the *Madcam1* promoter in Peyer’s patch (PP) HEC, peripheral lymph node (PLN) HEC, and PP capillary EC (CapEC). Blue highlight denotes site of conserved CE. (B) Sequence of the CE in indicated species. Conserved E-box (HEY1 binding site), homeodomain (NKX2-3 binding site) and COUP-TFII binding sites are highlighted in red, purple and green respectively. (C) Sequence for the wildtype probe (CE) and mutant probes used for EMSA. CE_ΔE/N harbors mutations in the HEY1 and NKX2-3 binding site; CE_ΔC_AB harbors mutations in the COUP-TFII “A” and “B” sites. Red, purple and green bars denote the HEY1, NKX2-3 and COUP-TFII binding sites, respectively. (D) EMSA showing migration of recombinant HEY1, NKX2-3 and COUP-TFII-GST bound to the CE probe (red arrows), supershifted by antibodies specific to the corresponding transcription factor (red stars). Cold competitor probe outcompeted the CE probe (lanes 4, 9, 14. Mutated probes lacking the HEY1 and NKX2-3 sites (CE_ΔE/N, lanes 5 and 10) or the COUP-TFII sites (CE_ΔC_AB, lane 15) failed to bind the respective TFs.

To test for direct binding of the TFs at the *Madcam1* CE, we performed electromobility shift assays (EMSA) using recombinant proteins and a labeled *Madcam1* CE probe (Figure 1C and D). Binding of HEY1, NKX2-3 and COUP-TFII to the *Madcam1* CE probe was confirmed by supershift of the TF:CE complexes with respective anti-TF antibodies (Figure 1D, lanes 3, 8 and 13). NKX2-3:CE binding was abolished by a mutation of the homeodomain site (Figure 1D, lane 10); and HEY1:CE binding was abolished by a mutation of the E-box motif (Figure 1D, lane 5). In addition, COUP-TFII:CE binding was blocked by mutation of the two COUP-TFII binding sites (Figure 1D; lane 15). These results confirm that the identified motifs mediate TF:CE binding. Moreover, as predicted, HEY1 dose-dependently inhibited NKX2-3 binding to the “A” site, indicating that binding to the overlapping motifs is competitive (Figure S2).

### The *Madcam1* CE confers negative regulation by Notch signaling via Hey1

To assess the function of the composite motif, we generated luciferase (LUC) reporter constructs driven by the proximal NF-κB site with or without the *Madcam1* CE, and transfected the constructs into bEnd.3 endothelial cells, a well-characterized mouse endothelial cell line that supports TNFα-induced *Madcam1* expression (Ogawa et al., 2005; Sikorski et al., 1993) (Figure 2A; sequence information in Supplemental Table S1). The proximal NF-κB site by itself drove robust TNFα-responsive luciferase expression. However, promoter activity was dramatically inhibited by inclusion of the CE (Figure 2A), suggesting that the CE engages inhibitory signals active in the cultured endothelial cells.

**Figure 2.**
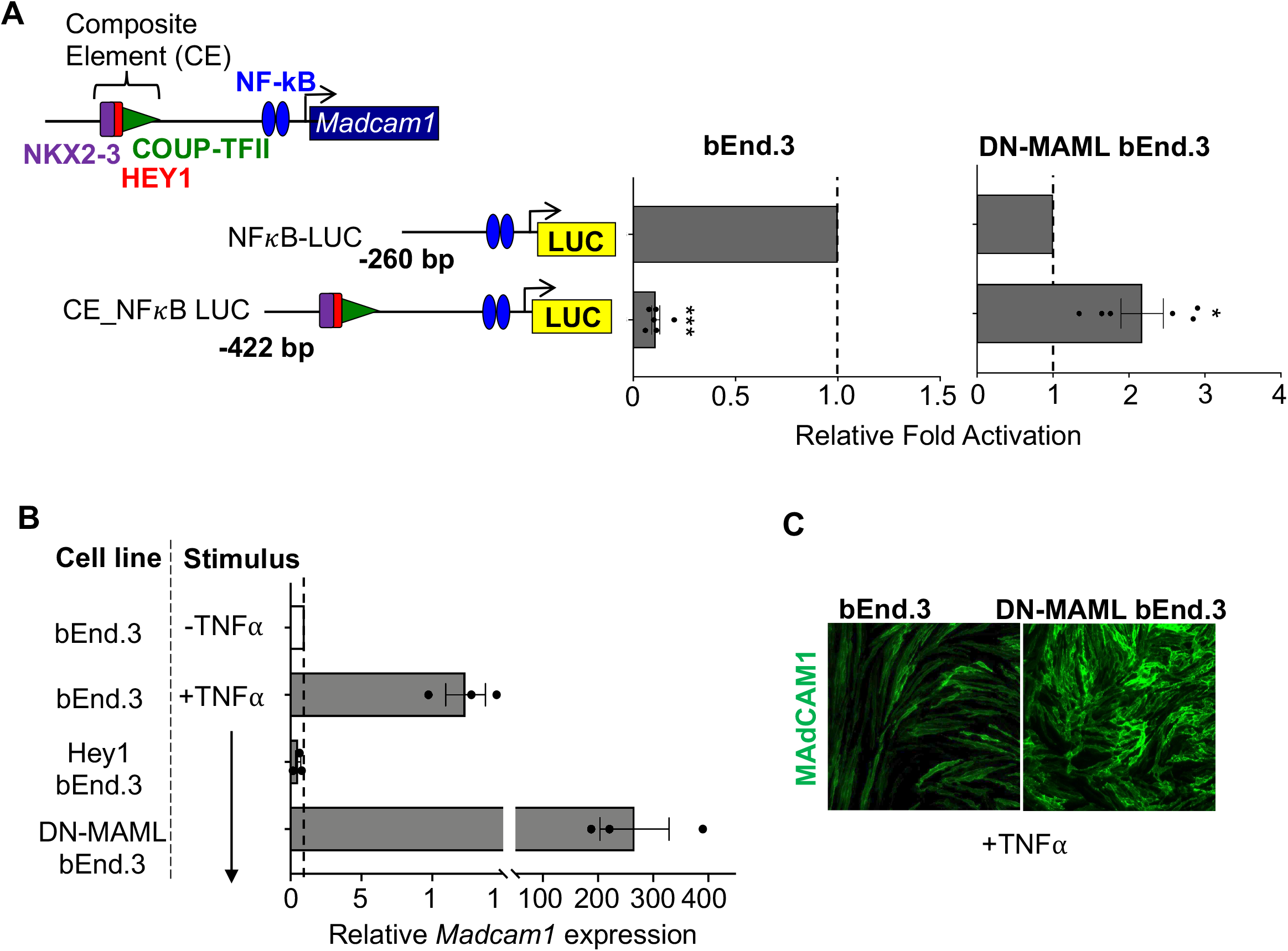
Notch signaling inhibits MAdCAM1 expression through the CE. (A) Luciferase reporter activity driven by the *Madcam1* promoter with (CE_NF**κ**B-LUC) or without (NF**κ**B-LUC) the CE. Data are from three biological replicates, each with two technical replicates for each condition. CE_NF**κ**B-LUC activity is normalized to the activity of NF**κ**B-LUC. ***: p-value <0.005, *: p-value <0.05, two tailed T-test, paired. (B) Endogenous *Madcam1* expression in bEnd.3 cells or bEnd.3 cells stably transduced with *DN-MAML* or *Hey1*, evaluated by real-time PCR. Results are normalized to basal *Madcam1* in unstimulated bEnd.3 cells. Data are from three biological replicates with SEM. (C) Immunofluorescence staining showing MAdCAM1 expression (green) in TNF□-stimulated bEnd.3 vs DN-MAML bEnd.3 cells. Except as indicated (white bars), transfected and control endothelial cells in A-C were stimulated with TNFα for 16-36 hrs prior to assay.

We reasoned that E-box binding transcriptional repressors, which are induced by Notch signaling, might mediate this transcriptional inhibition. To assess the function of the CE in the absence of Notch signaling, we used bEnd.3 endothelial cells with stable expression of dominant negative *MAML (DN-MAML)*, a mutant *Mastermind-like 1* gene which potently inhibits endogenous Notch-dependent expression of HEY1 and HES1 (Maillard et al., 2004). In contrast to its inhibitory effect in bEnd.3 EC, the CE-containing reporter construct significantly enhanced LUC activity in DN-MAML EC (Figure 2A). DN-MAML EC also showed increased basal and enhanced TNFα induction of endogenous *Madcam1* message (Figure 2B) and of surface MAdCAM1 protein expression (Figure 2C). Conversely, overexpression of *Hey1* suppressed MAdCAM1 induction at the transcript and protein level in bEnd.3 cells (Figure 2B and S3). Taken together, we conclude that Notch signaling, specifically the Notch downstream repressor HEY1, inhibits both CE activity and endogenous *Madcam1* induction in endothelial cells.

### NKX2-3 and COUP-TFII cooperatively enhance *Madcam1* expression in competition with HEY1

To determine the role of NKX2-3 and COUP-TFII binding at the *Madcam1* CE, we performed gain- and loss-of-function studies in bEnd.3 cells, which constitutively express low levels of *Nkx2-3* and *Nr2f2*. Overexpression of *Nkx2-3* in bEnd.3 cells enhanced TNFα-induced *Madcam1* expression (Figure 3A). Conversely, shRNAs targeting *Nkx2-3* or *Nr2f2* reduced *Madcam1* expression (Figure 3B). Thus, both NKX2-3 and COUP-TFII are required for endogenous *Madcam1* expression in endothelial cells. Anti-COUP-TFII chromatin immunoprecipitation (ChIP) in DN-MAML bEnd.3 cells showed enrichment for a DNA segment encompassing the *Madcam1* CE (Figure 3C), confirming *in vivo* COUP-TFII binding to the endogenous *Madcam1* promoter.

**Figure 3.**
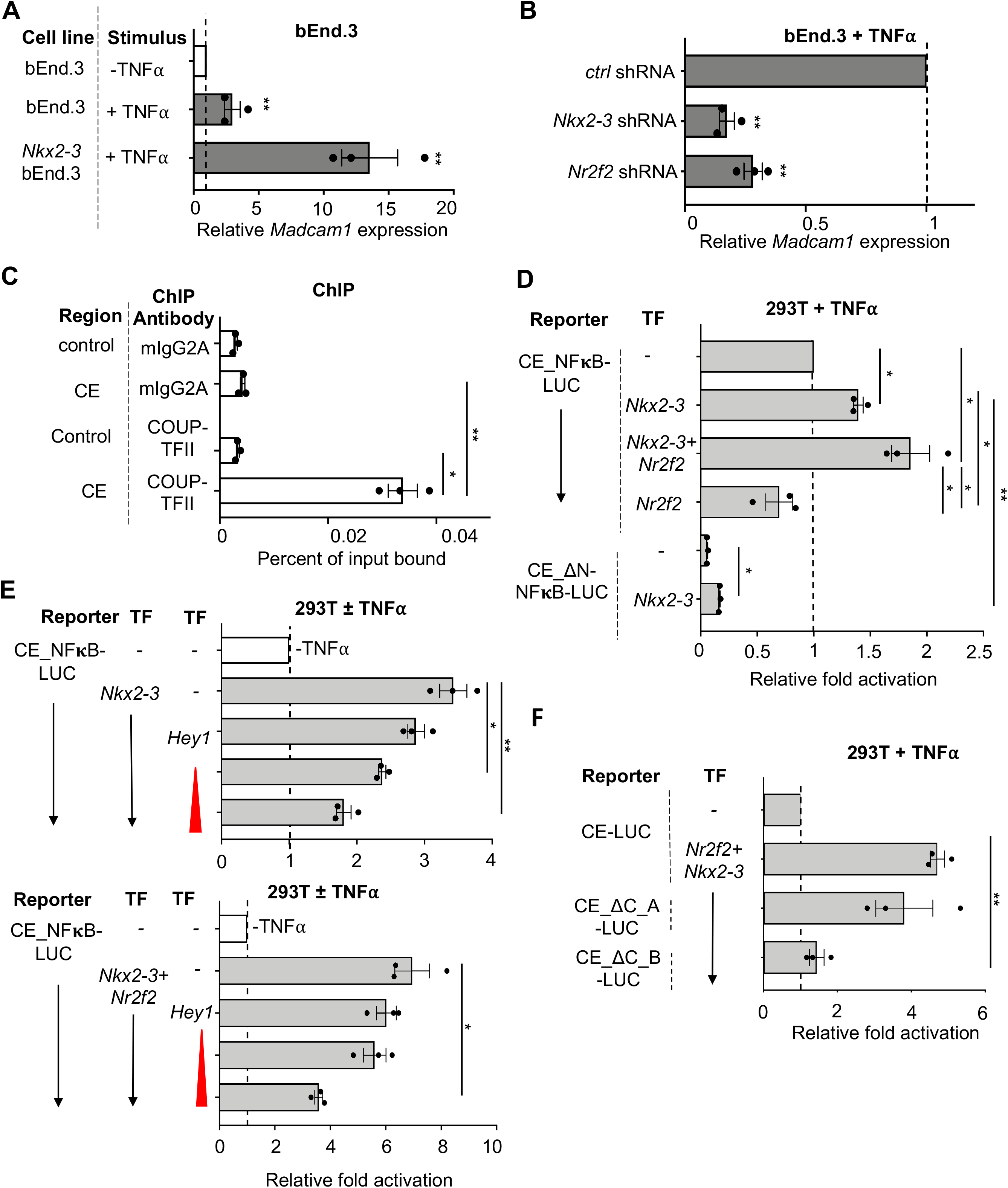
The *Madcam1* CE integrates NKX2-3 and COUP-TFII to activate transcription. (A) Induction of *Madcam1* expression in TNF□ stimulated bEnd.3 cells overexpressing *Nkx2-3*, evaluated by real-time PCR. Results are normalized to basal *Madcam1* levels in bEnd.3 cells. (B) Knockdown of *Nkx2-3* or *Nr2f2* by shRNA inhibits *Madcam1* expression in bEnd.3 cells. Data was normalized to β-actin then to expression in cells transduced with control shRNA. (C) Anti-COUP-TFII chromatin immunoprecipitation from nuclei of DN-MAML bEnd.3 cells enriches for the CE. Real-time PCR using primers for the CE was performed on the input and immunoprecipitated DNA fractions and results were shown as percent of input CE DNA. Precipitation of an irrelevant genomic control region without a COUP-TFII binding site was used as a negative control. Results are representative of three independent experiments. (D) *Nkx2-3* or cooperative *Nkx2-3* and *Nr2f2* enhance TNF□ stimulated luciferase reporter activity driven by CE-containing *Madcam1* promoter (CE_NF**κ**B-LUC) in 293T cells. Mutation of the homeodomain motif (in CE_ΔN-NF**κ**B-LUC) inhibits activity. Data normalized to CE_NF**κ**B-LUC activity. (E) HEY1 dose-dependently suppresses *Nkx2-3* (top) or cooperative *Nr2f2* and *Nkx2-3* (bottom) transcriptional activation from the CE-containing *Madcam1* promoter in 293T cells. Data normalized to basal reporter activity (white bars). Gray bars indicate TNF□ stimulation. (F) Effects of mutation of COUP-TFII “A” (CE_ΔC_A-LUC) or “B” (CE_ΔC_B-LUC) sites on cooperative *Nr2f2* and *Nkx2-3* activation of luciferase reporter in 293T cells. Results were normalized to control CE-containing reporter activity. Except as indicated (clear bars), in D-E cells were stimulated with TNF□ at the same time as transfection and assayed for LUC activity at 36 hours. All data are mean ± SEM of three independent experiments unless stated otherwise. ****: p-value <0.001, ***: p-value <0.005, **: p-value<0.01, *: p-value<0.05. Two tailed t-test, paired.

To evaluate the functional interplay of the TFs, we transiently transfected wild type or mutant CE-containing reporter constructs in 293T cells with or without co-transfecting *Nkx2-3, Nr2f2*, or *Hey1* and assessed luciferase induction by TNFα. *Nkx2-3* overexpression enhanced transcription from the wild type CE-containing reporter, whereas a mutation in the CE homeodomain motif (NKX2-3 binding site) abrogated this response, reducing reporter activity to well below the control without *Nkx2-3* overexpression (Figure 3D). On the other hand, overexpression of *Nr2f2* alone did not activate the reporter, but *Nr2f2* synergized with *Nkx2-3* to further promote enhancer activity (Figure 3D). Reporter activation by *Nkx2-3* or by *Nkx2-3* and *Nr2f2* was inhibited dose-dependently by *Hey1* (Figure 3E).

To address the role of the two COUP-TFII sites within the CE, we utilized a reporter consisting of tandem CE motifs without proximal *Madcam1* promotor sequences. Mutational analyses showed that activation of reporter expression in 293T cells by *Nr2f2* and *Nkx2-3* required the COUP-TFII “B” binding site, whereas mutation of the COUP-TFII “A” site had little or no effect (Figure 3F), suggesting recruitment of an activating COUP-TFII:NKX2-3 complex on the CE. We conclude that the *Madcam1* CE integrates cooperative transcriptional activation by *Nkx2-3* and *Nr2f2*, in competition with the Notch downstream repressor *Hey1*.

Although the spacing and sequence of candidate TF binding sites in the human *MADCAM1* CE differ from those in the mouse (Figure 1, S1), reporter assays confirmed that the CE regulatory functions are conserved: Basal as well as TNFα-driven human *MADCAM1* CE_ luciferase reporter activity was upregulated by transfection of *NKX2-3*, and cooperatively enhanced by co-transfection of *NKX2-3* and *NR2F2* (Figure S4A). Excess *NR2F2* reduced reporter activity, potentially reflecting competitive displacement of NKX2-3 by COUP-TFII at the “A” site. Consistent with the mouse data (Figure 3E), over-expression of *HEY1* antagonized *NKX2-3* driven reporter activity; and was also effective but less efficient at inhibiting the combined response to *NKX2-3* and *NR2F2* (Figure S4B). Tandem *MADCAM1* CE sequences were sufficient to confer reporter activity from a minimal promoter in response to *NKX2-3* and *NR2F2*, as well as negative regulation by *HEY1* (Figure S4C). These results support functional conservation of the regulatory interplay of NKX2-3, COUP-TFII and HEY1 for combinatorial control of *MADCAM1* transcription.

### NKX2-3 and COUP-TFII form DNA- and tinman-domain dependent heterodimers

Coactivation of the *Madcam1* CE-containing reporter by COUP-TFII and NKX2-3 as well as enrichment of *Madcam1* CE from COUP-TFII ChIP suggested coordinate binding of these TFs to the CE. To test this hypothesis, we used EMSA to assess interaction of NKX2-3 and COUP-TFII on the native and mutant CE probes (Figure 4A). Co-incubation of NKX2-3 and COUP-TFII with native CE yielded a band distinct from those formed by CE bound to NKX2-3 or COUP-TFII alone (Figure 4B, left, blue arrow). This band was intensified by mutation of the COUP-TFII “A” site (Figure 4B, right, lane 8), possibly reflecting competition of COUP-TFII and NKX2-3 for “A” site binding. Supershift with an NKX2-3 antibody confirmed NKX2-3 participation in the complex (Figure 4B, right, red star), whereas the control goat IgG antibody did not shift the protein:DNA complex (Figure 4B, right, lane 10). When we abolished both the NKX2-3 and COUP-TFII “A” binding sites, NKX2-3 no longer bound to the probe (Figure 4C, lane 2); but it retained the ability to bind together with COUP-TFII to form an NKX2-3:COUP-TFII:DNA complex (Figure 4C, lane 3, blue arrow; lane 5, red star). Formation of the heterotrimeric NKX2-3:COUP-TFII:DNA complex even in the absence of a canonical NKX2-3 binding site suggests that COUP-TFII, bound to DNA through a minimal motif, can initiate formation of the gene regulatory complex through protein-protein interactions which could then be stabilized by the adjacent homeodomain motif.

**Figure 4.**
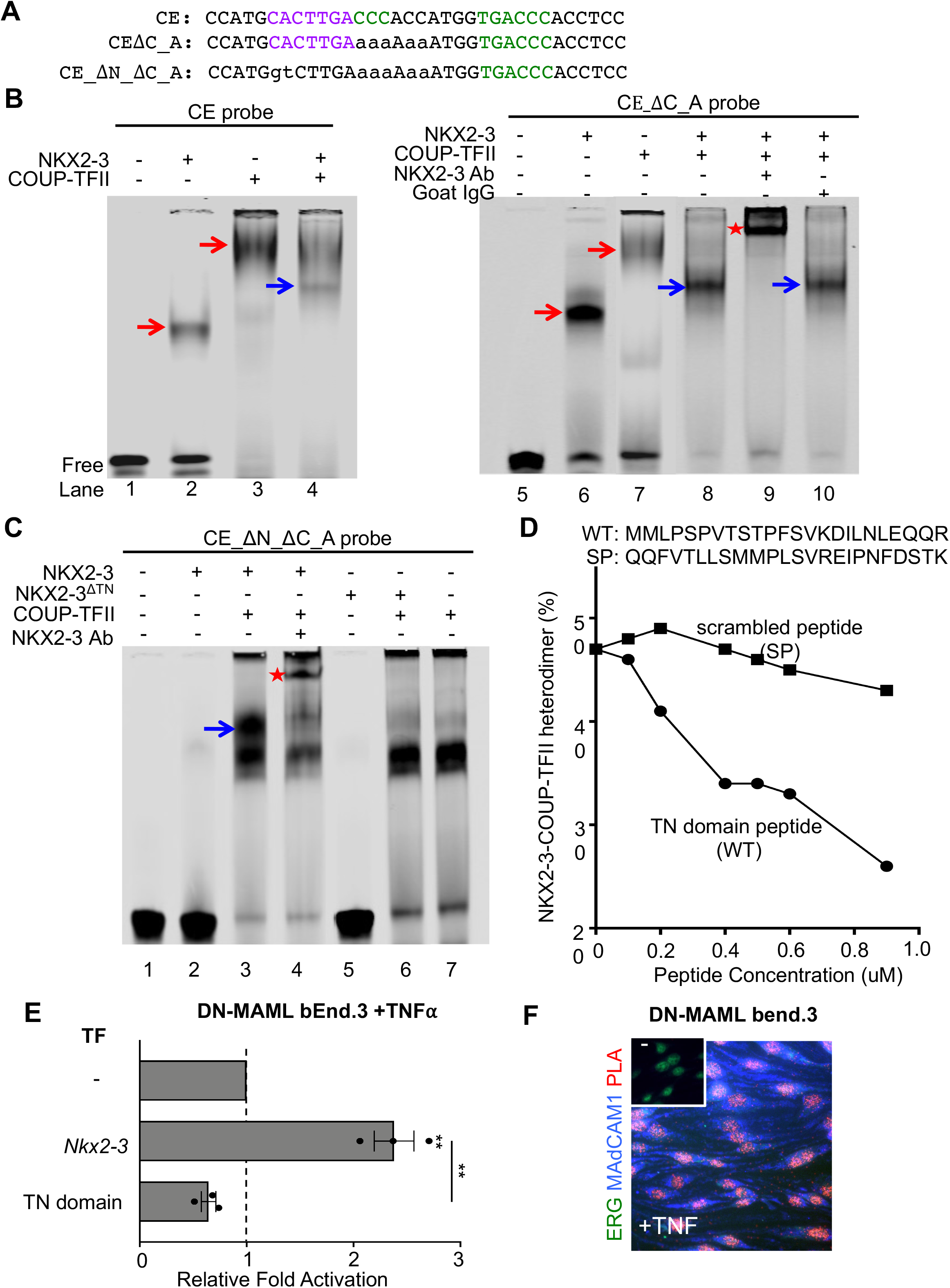
NKX2-3 and COUP-TFII heterodimerize via the tinman (TN) domain to activate MAdCAM1. (A) Sequence of wildtype and mutant CE probes for EMSA. (B) EMSA showing migration of recombinant NKX2-3 and COUP-TFII bound to the CE probe (left) and to CE_ΔC_A probe that lacks the COUP-TFII “A” site (right) (red arrows). Co-incubation of NKX2-3 and COUP-TFII yields a distinct band (blue arrows) reflecting an NKX2-3:COUP-TFII DNA complex that is supershifted by anti-NKX2-3 antibody but not by control antibody. (C) EMSA of protein complexes formed by incubation of the indicated TFs with a mutant CE probe lacking the NKX2-3 and the COUP-TFII “A” binding sites (CE_ΔN_ΔC_A). The probe binds COUP-TFII (lane 7) and not NKX2-3 (lane 2), but seeds a COUP-TFII:NKX2-3:DNA complex (blue arrow) when co-incubated with both TFs. The heterodimer-DNA complex is supershifted by anti-NKX2-3 antibody (red star). NKX2-3^ΔTN^ failed to heterodimerize with COUP-TFII (lane 6). (D) Disruption of the NKX2-3:COUP-TFII heterodimer-DNA complex by a competing NKX2-3 TN domain peptide (WT) or by a scrambled peptide (SP). Data is expressed as percent of probe bound by NKX2-3:COUP-TFII heterodimer. (E) *Madcam1* expression in TNFα-treated DN-MAML bEnd.3 cells stably overexpressing *Nkx2-3* or the *Nkx2-3* TN domain, as evaluated by real-time PCR. Values are mean ± SEM of three biological replicates. Two tailed t-test, paired. **: p-value <0.01. (F) Proximity ligation assay of NKX2-3 and COUP-TFII in TNFα-treated DN-MAML bEnd.3 cells stained for MAdCAM1 (blue) and ERG (green), with the proximity ligation signal shown in red. Inset shows a negative control with isotype matched primary antibodies.

NKX2-3 contains an N-terminal tinman (TN) domain, conserved in members of the NKX2 family. Recombinant NKX2-3 lacking the TN domain failed to dimerize with COUP-TFII on CE (Figure 4C, lane 6). A peptide encompassing the TN domain inhibited NKX2-3:COUP-TFII heterodimerization, while a scrambled peptide had no effect (Figure 4D); and overexpression of the TN domain peptide suppressed basal *Madcam1* and abrogated the enhanced *Madcam1* induction seen in DN-MAML bENd.3 cells transfected with NKX2-3 (Figure 4E). Thus, NKX2-3:COUP-TFII complex formation and function are dependent on physical interactions involving the NKX2-3 TN domain. Proximity ligation assay (PLA) confirmed the physical association of NKX2-3 and COUP-TFII within the nuclei of EC expressing MAdCAM1 (Figure 4F).

### Identification of a conserved *St6gal1* NCCE

*St6gal1*, encoding the β-galactoside α-2,6-sialyltransferase 1, confers expression of the B cell-recruiting glyco-addressin BMAd, and like *Madcam1* is selectively expressed by intestinal HEV. We identified a conserved NKX-COUP-TFII composite element (NCCE) within an enhancer region in the first intron of *St6gal1* (Figure 5A), suggesting a common genomic address code might regulate the expression of both mucosal addressins. EMSA showed that NKX2-3 and COUP-TFII each bound the native *St6gal1* NCCE (Figure 5B). Binding to the labeled CE was inhibited by cold wild type probe but not by mutant probes lacking the cognate homeodomain or COUP-TFII motifs (Figure S5). Co-incubation with NKX2-3 and COUP-TFII produced a distinct complex (Figure 5B, blue arrow), indicative of heterodimer formation as seen with the *Madcam1* CE (Figure 4B). Endogenous expression of *St6gal1* in DN-MAML bEnd.3 cells was inhibited by shRNA knockdown of *Nr2f2* or *Nkx2*-3 (Figure 5C) and conversely upregulated by overexpressing *Nkx2-3* (Figure 5D). *St6gal1* was effectively suppressed by overexpression of the NKX2-3 tinman domain (Figure 5D), supporting the requirement for both TFs and for TN-dependent interactions for optimal *St6gal1* expression in EC. Tandem *St6gal1* NCCE sequences activated a minimal luciferase reporter in response to transfected COUP-TFII or NKX2-3, and co-transfection of both TFs further enhanced activation (Figure 5E, gray bars). Mutation of the COUP-TFII site suppressed the cooperative response (Figure 5E, white bars). Thus the intronic NCCE in *St6gal1*, like that in the *Madcam1* promoter, is sufficient to confer combinatorial activation of gene expression by NKX2-3 and COUP-TFII.

**Figure 5.**
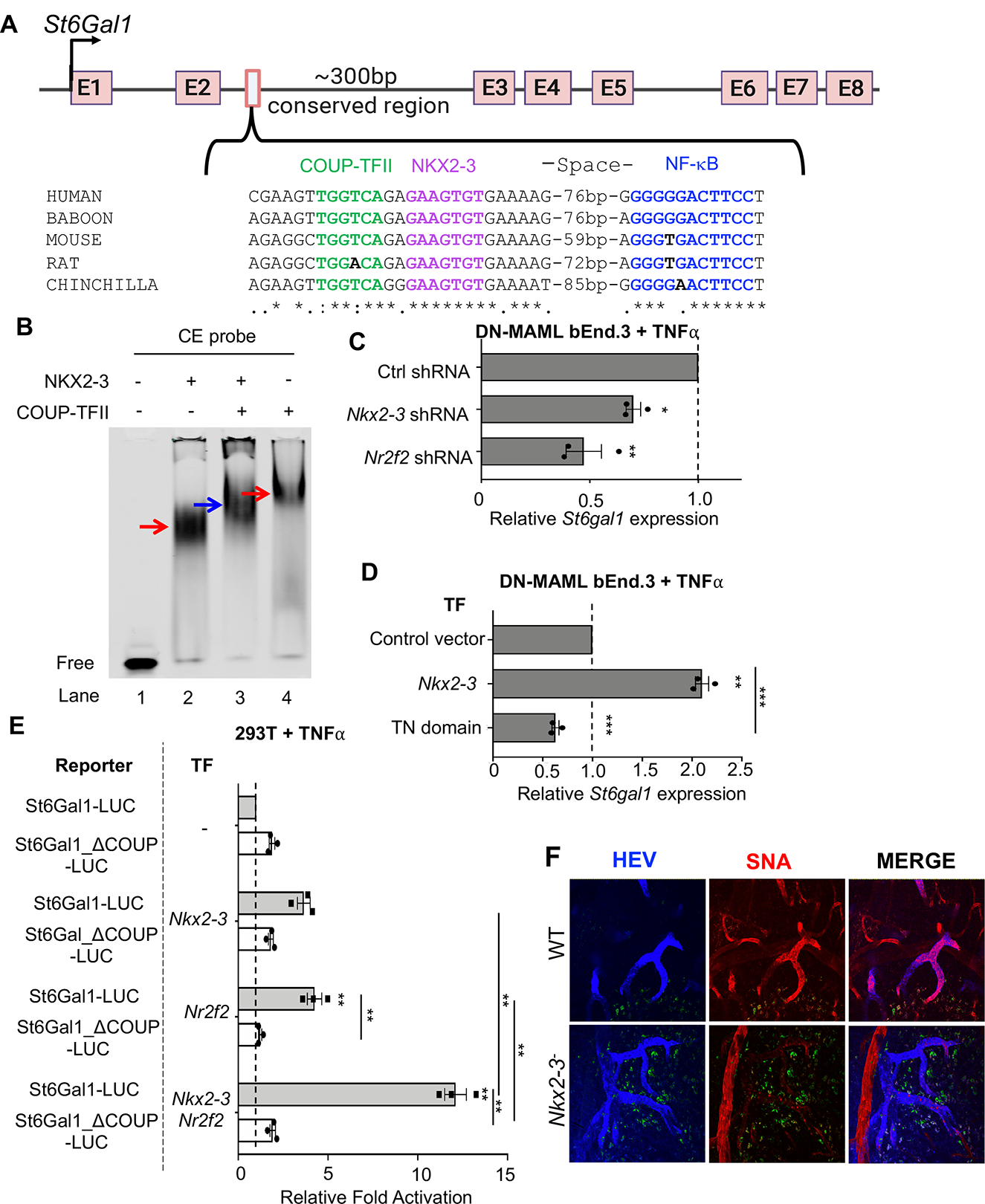
NKX2-3 regulates *St6gal1* expression via a conserved NCCE. (A) Schematic of the *St6gal1* gene and an NCCE in the second intron. (B) EMSA showing migration of NKX2-3, COUP-TFII-GST (red arrows) and NKX2-3: COUP-TFII heterodimer (blue arrow) bound to an *St6gal1* CE probe. (C) *St6gal1* expression in DN-MAML bEnd.3 cells stably transfected with *Nkx2-3* or *Nr2f2* shRNA, evaluated by real-time PCR. (D) *St6gal1* expression in DN-MAML bEnd.3 cells stably transfected with NKX2-3 or the NKX2-3 TN domain. Data shown as mean ± SEM of three biological replicates, each with 3 technical replicates per condition. (E) Activity of luciferase reporter driven by *St6gal1* NCCE-containing promoter (St6Gal1-LUC) or *St6gal1* NCCE-containing promoter with mutated COUP-TFII binding sequence (St6Gal1_ΔCoup-LUC), when co-transfected with *Nr2f2* and/or *Nkx2-3* in 293T cells. Results are shown as mean ± SEM from three independent transfections. Two tailed t-test, paired. *: p-value <0.05; **: p-value <0.01; ***: p-value <0.005. (F) Immunofluorescence of alpha-2,6-sialic acid binding lectin SNA staining in PP HEV in WT vs NKX2-3 deficient mice (red). HEV are marked by AF450-labeled anti-addressin antibodies MECA79, MECA89 and MECA367. Autofluorescence is shown in green.

### NKX2-3 is required for BMAd expression in Peyer’s patch HEV

The identification of the *St6gal1* NCCE suggested that NKX2-3 might drive BMAd expression in gut venules. To address this hypothesis, we used the *Sambucus nigra* lectin (SNA) to assess vascular display of α-2,6-sialic acid glycotopes. SNA stained PP HEV intensely in WT mice, but reactivity was abrogated in *NKX2-3^-/-^* mice (Figure 5F), indicating that NKX2-3 is required for *St6gal1*-dependent BMAd expression *in vivo*.

### Endothelial COUP-TFII overexpression induces ectopic *Madcam1* and *St6gal1* in gut capillaries

To evaluate the role of COUP-TFII in regulation of the mucosal addressins *in vivo*, we generated mice with inducible EC-specific overexpression (EC-iCOUP^OE^) or deficiency (EC-iCOUP^KO^) by crossing Cadherin 5-cre^ERT2^ mice (Sorensen et al., 2009) to lox stop lox Rosa COUP-TFII (COUP^KO^) (Takamoto et al., 2005) or to Nr2f2^tm2Tsa^ (COUP^OE^) mice (Li et al., 2009), respectively (Figure 6A).

**Figure 6.**
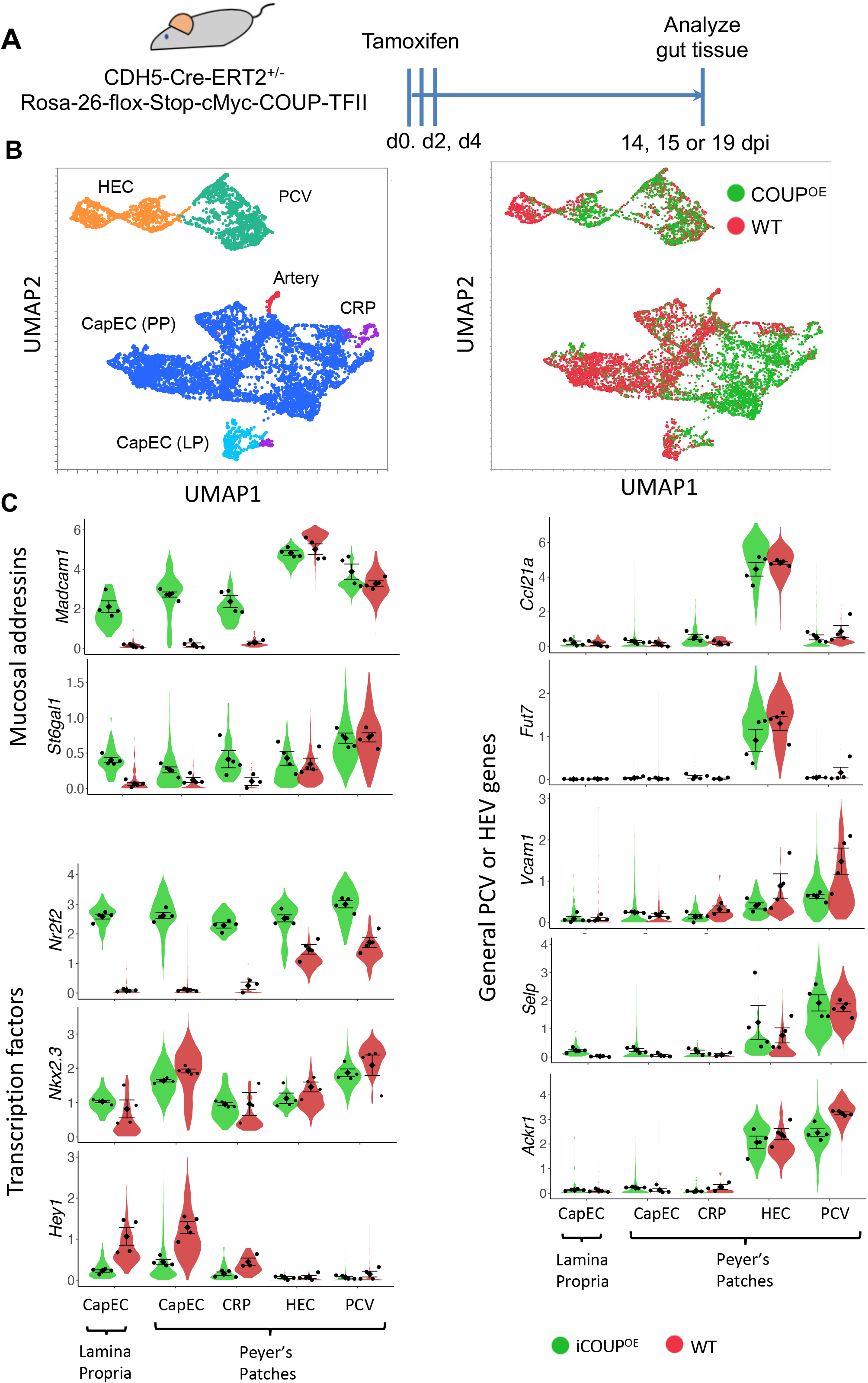
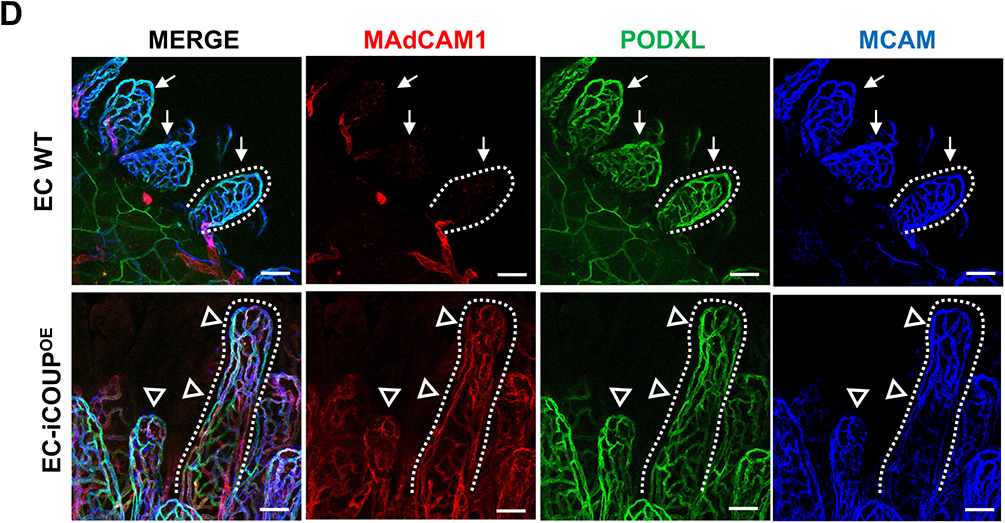
Pan-endothelial COUP-TFII expression induces ectopic *Madcam1* and *St6gal1* in intestinal capillaries. (A) Experimental timeline for induction of COUP-TFII in EC. (B) UMAP plot of PP BEC colored by EC subset (left) or by genotype (right). (C) Violin plots showing expression of select genes in EC-iCOUP^OE^ vs control BEC subsets. Dots represent mean gene expression of separate cohorts (male and female cohorts from two independent experiments), and are presented with SEM. (D) Confocal imaging showing MAdCAM1 expression in lamina propria capillaries in EC-iCOUP^OE^ (arrowheads) but not control (arrows) mice. Scale bars: 50µm, EC stained by i.v. injection of directly conjugated antibodies. 20x, whole mount. See also Supplemental Figure 7 for MAdCAM1 extension into capillary segments in PP.

Pan-endothelial COUP-TFII overexpression was induced with tamoxifen, and 15 or 19 days later BEC were isolated from PP of male and female EC-iCOUP^OE^ mice and profiled by single cell RNA sequencing (scRNAseq) (Figure 6A). BEC from tamoxifen-treated Cre^-^ control and WT mice were also profiled. Male and female cohorts within the EC-iCOUP^OE^ and Cre^-^ cohorts were processed together and resolved post-sequencing. UMAP clustering separated the major capillary, arterial and venous EC populations (Figure 6B) as well as capillary resident angiogenic progenitors (CRP) (Brulois et al., 2020). Venous EC comprised *Madcam1*-high HEC and *Selp*-high, *Madcam1*-low PCV (Figure 6B and 6C). UMAP separated capillary phenotype EC characteristic of lymphoid tissues (the larger population) and putative lamina propria EC, the latter characterized by enrichment for genes for lipoprotein and cholesterol transport and for intestinal endothelial barrier formation. With the exception of mature *Gkn3*^+^ arterial EC (identifiable among captured cells only in the WT cohort), and of CRP (missing in the male WT mice), all subsets were represented by cells within both male and female mice from each condition.

CapEC in control cohorts lacked *Nr2f2* but expressed *Nkx2-3* (Figure 6C). Tamoxifen treatment induced *Nr2f2* expression in the majority of EC including the CapEC populations and CRP, and led to their upregulation of both *Madcam1* and *St6gal1* (Figure 6C). Rare CapEC within induced EC-iCOUP^OE^ mice that failed to express *Nr2f2* also failed to induce the addressin genes (not shown), even though they were *Nkx2-3^+^*, supporting the requirement for TF co-expression. Notably, the regulatory effect was selective for the mucosal addressins: *Nr2f2* overexpression did not induce or alter expression of genes encoding the non-intestinal or non-organotypic venous markers *Vcam1, Ackr1*, *Ccl21a*, or *Selp* (Figure 6C). Immunolabeling and confocal imaging confirmed widespread ectopic MAdCAM1 expression in capillaries in the gut lamina propria and in PP of EC-iCOUP^OE^ mice (Figure 6D and S7). Ectopic MAdCAM1 was observed in the capillary arcades of the small and large intestinal villi as early as 14 days after tamoxifen treatment (Figure 6D). Conversely, induced pan-endothelial COUP-TFII deficiency in EC-iCOUP^KO^ mice led to reduced MAdCAM1^+^ HEV in PP (Figure S8).

SNA-binding glycotopes *in vivo* are less restricted to HEV than MAdCAM1. In particular, many arteries and peripheral capillary segments arising from arteries in the intestinal lamina propria also bound the lectin in WT mice (Figure S6A, C and S9). However, in induced EC-iCOUP^OE^ mice SNA stained more extensively and decorated many of the CapEC that ectopically expressed MAdCAM1 in the mesh-like capillary network within lamina propria villi (Figure S9), consistent with the altered *St6gal1* expression pattern seen in EC-iCOUP^OE^ CapEC (Figure 6C).

### NCCE in the genome

The ability of NCCE to foster cooperative COUP-TFII:NKX protein interactions to activate gene expression led us to survey the genome for conserved NCCE motifs. In particular, we hypothesized that NCCE might be enriched in association with genes that drive embryonic heart morphogenesis and pancreatic beta cell specification, developmental events which are known to involve both NKX homeodomain factors and COUP-TFII (Anderson et al., 2009; Boutant et al., 2012; Doyle et al., 2007; Harvey, 1996; Lyons et al., 1995; McElhinney et al., 2003; Pereira et al., 1999; Wu et al., 2013; Wu et al., 2015). We screened conserved genomic regions for COUP-TFII motifs (GGTC(A/G)) with adjacent NKX HD motifs. We focused on NCCE with a 6-16 bp spacing between motif centers, which excluded overlap of the COUP-TFII and NKX sites and encompassed the range of spacings in NCCE we showed biochemically to seed the heterodimer. We identified 1954 genes nearest to these NCCE (NCCE+ genes; genomic information provided in Supplementary Table S4 and in a UCSC Genome Browser session (see Methods)). Since NKX and COUP-TFII motifs can regulate genes independently, as a control we identified genes lacking NCCE but with nearby conserved syntenic COUP-TFII and NKX HD motifs separated by larger spacings (30-60 bp), less likely to foster efficient DNA-dependent heterodimers (NCCE-genes). As additional controls, we identified genes with conserved NKX motifs 6-16 bp from scrambled motifs GTAC(G/C), AGTC(G/C), and TGGA(C/T).

While not all functional motifs need be conserved, and not all bioinformatically defined NCCE are likely to function *in vivo*, GO term analyses strongly support the physiologic relevance of the conserved NCCE: Compared to control genes, NCCE+ genes are associated, as hypothesized, with genes for morphogenesis of the heart and for pancreatic beta cell specification (Figure 7A). Moreover, NCCE are highly enriched in genes involved in cell specification and morphogenesis in multiple organ systems: neuron development and axonogenesis; morphogenesis of arteries, teeth, digestive tract, inner ear, cartilage, skeletal muscle, hind limb and joints (Figure 7B). Literature search identifies involvement or regulated expression of *Nkx* family members and *Nr2f2* in several of these processes, but their interplay has not been highlighted.

**Figure 7.**
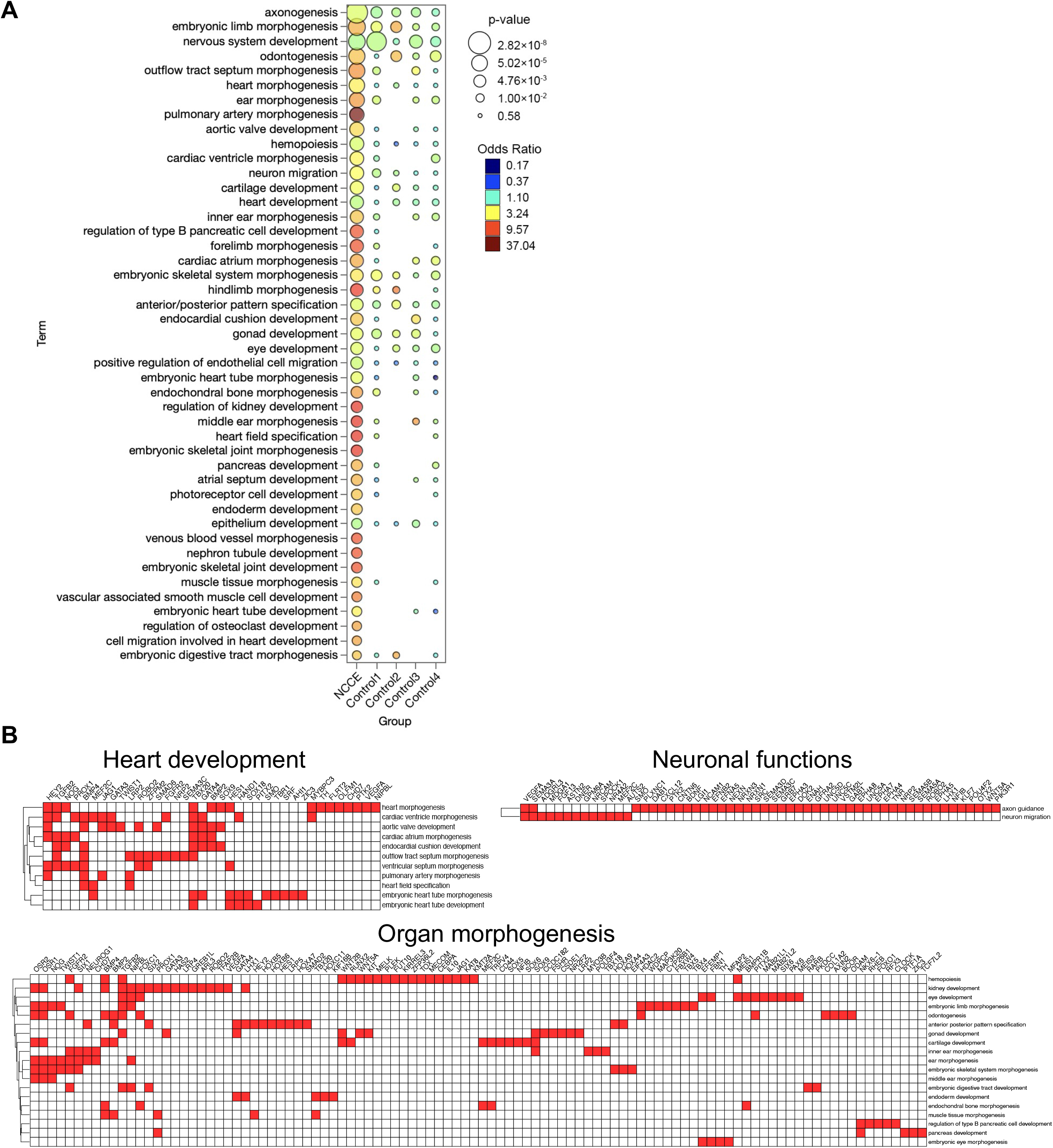
Genome-wide NCCE are associated with organ morphogenesis. (A) Select GO terms enriched in NCCE+ genes compared to controls. NCCE+ genes are defined as genes nearest to conserved NCCE, where the NCCE are defined as having a 6-16 bp spacing between the centers of a COUP-TFII and an NKX HD motif. Control 1: genes associated with conserved COUP-TFII and NKX HD motifs separated by larger spacings (30-60 bp). Controls 2-4: genes associated with conserved NKX motifs 6-16 bp from conserved scrambled motifs GTAC(G/C), AGTC(G/C), or TGGA(C/T), respectively. (B) NCCE+ genes in select pathways associated with heart development, neuronal functions, and organ morphogenesis.

To confirm the potential of candidate NCCE to facilitate cooperative interaction of COUP-TFII and NKX proteins, we carried out *in vitro* analyses of NCCE in regulatory regions of three NCCE+ genes, *Tbx5, Nrp2 and Etv2* (Figure S10). These genes were selected for the following reasons: 1) The NKX binding motifs within them are representative of the diversity of candidate NKX motifs identified informatically, 2) They are induced developmentally in cells that co-express *Nr2f2* and *Nkx* ((Harris and Black, 2010; Kanatani et al., 2015a; Schwach et al., 2017) or as shown here (Figure S11), and 3) Their expression and function in embryogenesis depend on both NKX family HD proteins and on COUP-TFII. NKX2-1 and COUP-TFII coordinate *Nrp2-*dependent development and migration of GABAnergic neurons (Kanatani et al., 2015b). NKX2-5 with COUP-TFII regulates *Tbx5* and *Etv2* in cardiac development (Ferdous et al., 2009a; Nobrega-Pereira et al., 2008; Wu et al., 2013; Wu et al., 2015). (Moreover, the NKX motif in the *Etv2* NCCE has been previously shown to bind NKX2-5 (Ferdous *et al*., 2009a; He et al., 2011; Nobrega-Pereira *et al*., 2008)).) EMSA confirmed that the conserved *Tbx5* and *Etv2* NCCE support formation of COUP-TFII:NKX2-5 heterodimers (Figure S12A, B and D), and the *Nrp2* NCCE seeds formation of COUP-TFII:NKX2-1 heterodimers (Figure S12A and F). Moreover, tandem NCCE sequences from each of these genes integrated COUP-TFII and NKX2-5 or NKX2-1 to cooperatively activate reporter expression from a minimal promoter (Figures S12C, E and G, respectively). Taken together, these results define a library of conserved genomic NCCE, confirm that diverse NCCE can seed functional NKX-COUP-TFII heterodimers, and implicate NCCE in cell specification and morphogenesis in development.

## DISCUSSION

Lymphocyte recruitment to the intestines and GALT is regulated by two intestinal vascular addressins, MAdCAM1, an Ig family member for lymphocyte integrin α_4_β_7_ (Berlin et al., 1993), and BMAd, a B cell-specific addressin comprising *St6gal1-*dependent α-2,6-sialic acid glycotopes for recognition by B cell CD22 (Lee *et al*., 2014a). Here we have identified an NKX-COUP-TFII composite element (NCCE) that functions as a genomic address code for the organ- and venule-specific expression of these adhesion molecules. We show that the NCCE binds developmentally programmed TFs NKX2-3 and COUP-TFII, which form heterodimers that activate transcription of *Madcam1* and *St6gal1* in concert with NF-κB signals. The NKX tinman domain, which is required for the interaction of NKX2-3 with COUP-TFII, is known to recruit a Groucho co-repressor complex that mediates transcriptional repression (Muhr et al., 2001). Association with COUP-TFII may mask the tinman binding site for Groucho, or alter tinman protein partners to instead activate transcription. We show that NKX2-3 is required for *St6gal1*-dependent BMAd expression by intestinal Peyer’s patch HEV, as shown previously for MAdCAM1 (Pabst *et al*., 2000). Moreover, transgenic endothelial expression of COUP-TFII is sufficient to induce ectopic expression of *Madcam1* and *St6gal1* in intestinal capillaries which express NKX2-3. Finally, we show that the *Madcam1* CE comprises an additional control element that binds capillary/artery-expressed repressors downstream of Notch signaling, reinforcing the requirement for COUP-TFII and NKX2-3 and the selectivity of MAdCAM1 for venules. Together the results show that the conserved CE integrate master intestinal, venous and capillary/arterial TFs for tight control of addressins that regulate lymphocyte homing into the intestines and their associated lymphoid tissues.

NKX2-3 is expressed in the developing spleen, midgut, hindgut, and pharyngeal endoderm and contributes to development of intestinal vascular and stromal cells (Pabst *et al*., 2000; Pabst et al., 1997; Pabst et al., 1999; Vojkovics et al., 2018; Wang *et al*., 2000). In the adult it is retained by both venous and capillary/arterial EC within the intestines (Lee *et al*., 2014a), but not by EC in peripheral tissues or lung. Splenic marginal sinus endothelial cells express MAdCAM1, and NKX2-3 deficient mice lack MAdCAM1 expression in spleen as well as in the gut, consistent with a fundamental requirement for endothelial NKX2-3 expression in the adult (Kellermayer et al., 2014). Interestingly, in *Nkx2-3^-/-^* mice both GALT HEV and spleen sinus endothelial cells assume a PLN HEV-like phenotype with expression of the peripheral lymph node vascular addressin (PNAd) (Czömpöly *et al*., 2011) (Kellermayer *et al*., 2014), suggesting that NKX2-3 drives organotypic EC specialization through combined activation of intestinal EC gene expression and repression of peripheral (non-intestinal) vascular phenotypes. Our finding that NKX2-3 deficiency abrogates PP HEV BMAd expression is consistent with the transdifferentiation towards a peripheral HEV phenotype.

In contrast to its restricted expression in the adult, MAdCAM1 is widely displayed by endothelial cells during fetal and early neonatal life (Mebius et al., 1996). In neonates it is expressed by HEV in the peripheral lymph nodes as well as in developing GALT (Mebius *et al*., 1996) reviewed in (Ager, 2017). MAdCAM1 is progressively downregulated in PLN HEC, with concurrent upregulation of PNAd by 4-6 weeks of age. NKX2-3 deficiency has no effect on perinatal endothelial MAdCAM1, nor on the postnatal induction of PNAd in PLN, instead allowing emergence of peripheral characteristics in intestinal HEV with loss of MAdCAM1 in adults (Kellermayer *et al*., 2014). While our results provide an explanation for selective MAdCAM1 expression by intestinal venules in the adult, mechanisms underlying the NKX2-3-independent *Madcam1* expression in neonates and the NKX2-3-dependent suppression of PNAd in adult HEV remain to be defined.

Genome wide association studies implicate NKX2-3 variants in susceptibility to both Crohn’s disease and ulcerative colitis (Lu et al., 2014). While the basis for these associations remains unclear, as confirmed here *NKX2-3* enhances endothelial expression of MAdCAM1, and allelic studies show that *NKX2-3* risk haplotypes are overexpressed compared to non-risk alleles in the colon (Arai et al., 2011). Moreover, *NKX2-3* and *MADCAM1* are upregulated in IBD patients (Arai *et al*., 2011; Briskin *et al*., 1997), and MAdCAM1 is involved in the pathogenesis of inflammatory bowel disease as indicated by disease suppression by antibodies to MAdCAM1 and its lymphocyte receptor integrin α4β7 (Habtezion et al., 2016; Lamb et al., 2018). Together the available results could support a proposed link between variant *NKX2-3* and endothelial cell regulation of immune cell traffic and adhesion.

The other key component of the NCCE control complex, COUP-TFII, is an evolutionarily conserved orphan nuclear receptor that regulates diverse biological processes including angiogenesis, cardiac chamber development, neuronal subtype specification, circadian rhythm and metabolism (Park et al., 2003; Pereira et al., 1999). COUP-TFII acts as a transcriptional activator or repressor in a gene- and cell-type-dependent manner. During embryonic endothelial fate specification, COUP-TFII supports venous differentiation by repressing Notch signaling and arterial differentiation (You et al., 2005) and its expression is maintained in most venular endothelial cells *in situ* in the adult (Brulois *et al*., 2020; Kalucka et al., 2020). COUP-TFII forms stable homodimers in solution that bind to direct repeats (DR) of the COUP-TFII half site motif with variable spacing between repeats. Depending on the spacing, these DR sequences can also bind heterodimers of COUP-TFII with the nuclear hormone receptor RXR (Perlmann et al., 1996) (Qin et al., 2014). COUP-TFII is classically repressive in these settings (Butler and Parker, 1995) (Qin *et al*., 2014). COUP-TFII can also regulate gene expression positively in complex with other DNA-binding TFs (e.g. SP1, HNF4 or PROX1) independently of cognate COUP-TFII DNA binding sites (Mehanovic et al., 2019) (Ktistaki and Talianidis, 1997) (Lee et al., 2009; Yamazaki et al., 2009)): These regulatory complexes are not associated with conserved genomic sites for COUP-TFII-DNA interaction, and thus appear to be driven by protein-protein recognition rather than by DNA codes. In contrast, while our gel shift assays show that DNA-bound COUP-TFII can recruit NKX2-3 in the absence of a canonical NKX motif, this interaction is not sufficient to drive reporter expression in our systems: Activating NKX-COUP-TFII heterodimers may require stabilization by canonical NCCE in which both TFs bind their cognate motifs. Certainly, the biological significance of the *Madcam1* and *St6gal1* NCCE studied here is supported by their evolutionary conservation.

Enrichment of conserved NCCE in genes associated with embryonic organ morphogenesis suggests that COUP-TFII:NKX heterodimer-seeding motifs have significant roles beyond regulation of the addressins. COUP-TFII and the NKX homeodomain TF family often impact common cell fate decisions in development. NKX2-5, the vertebrate homolog of Drosophila ‘tinman’, establishes the embryonic heart field (Lyons et al., 1995), and COUP-TFII drives atrial fate determination within the heart field by inducing TBX5, a master transcription factor for atrial development (Wu *et al*., 2013) (Wu et al., 2016). Our results show that genes involved in heart field specification and in atrial development are enriched in NCCE+ genes (Figure 7B and C) including *Tbx5*, whose intronic NCCE we characterize biochemically and functionally (Figure S12A-C). COUP-TFII and NKX2-5 also contribute to and are independently required for pro-epicardial and endocardial development, and NCCE+ genes are enriched in genes associated with endocardial cushion morphogenesis. NKX2-5 transactivates endothelial expression of the Ets family member *Etv2* for COUP-TFII-regulated endocardial specification (Ciofani et al., 2012; Ferdous et al., 2009b; Lin et al., 2012), and we identify a conserved NCCE in the *Etv2* promoter that drives cooperative activation of a reporter gene by NKX2-5 and COUP-TFII (Figure S12A, D and E). NCCE+ genes are enriched in genes associated with neuron migration which include *Nrp2.* NKX2-1 and COUP-TFII are coordinately expressed and induce *Nrp2* in E13.5 GABAergic neurons, triggering their migration toward the amygdala or neocortex (Elias et al., 2008; Kanatani *et al*., 2015a; Tang et al., 2012). We show that an evolutionarily conserved NCCE in the *Nrp2* promoter supports NKX2-1/COUP-TFII interaction and reporter activation (Figure S12A, F and G). NCCE are also enriched among genes implicated in inner ear morphogenesis; in odontogenesis (which involves COUP-TFII and NKX2-3 (Han et al., 2018; Hur et al., 2015); and pancreatic beta cell fate specification (regulated by COUP-TFII and NKX2-2 (Boutant et al., 2012; Gutierrez et al., 2017; Papizan et al., 2011). In contrast, genes in which NKX and COUP-TFII sites are all separated by 30-60 bp show significantly less enrichment in NCCE-associated GO terms for morphogenesis or organogenesis, and/or are enriched in distinct embryonic and developmental processes. Thus, while the physiologic roles of the individual NCCE presented here as a resource remain to be determined, taken together the results suggest that NCCE contribute to diverse cell fate decisions and differentiation events in vertebrate organogenesis.

The study has limitations. Additional investigations will be required to fully explore the functional capabilities and physiologic roles of the NCCE identified here, including the *Madcam1* and *St6gal1* NCCE. Unanswered questions include whether other homeodomain and/or nuclear hormone receptors can interact competitively at the GGTCA and HD motifs, regulating or replacing NKX-COUP-TFII heterodimer-dependent gene expression and functions; whether NCCE and NKX-COUP-TFII heterodimers can recruit repressive as well as activating regulatory complexes; and how NKX-COUP-TFII heterodimers (and these potential additional mechanisms at NCCE) affect gene expression *in vivo* in different cell types and models. Targeted *in situ* mutagenesis of HD and COUP-TFII sites within NCCE in animal models could further illuminate the complex roles of NCCE and, in combination with co-expression/knockdown studies of additional TF in endothelial models, could provide additional insights into the intersection of mechanisms at NCCE that control physiologic gene expression. Second, the importance of spacing of COUP-TFII and HD motifs in NCCE remains incompletely characterized. We identify NCCE with gaps of 6-16 bp that can seed cooperative NKX-COUP-TFII binding and activate gene expression, and show that conserved NCCE with gaps in this range are enriched in functional gene associations; but understanding how the efficiency of heterodimer formation declines beyond this range, and whether NKX-COUP-TFII interactions can help bring distance sequences together, will require future biochemical and functional studies. Third, additional studies are required to understand the structural basis for heterodimer formation, and specifically how the tinman domain mediates the interaction. Fourth, the atlas of evolutionarily conserved NCCE we provide is necessarily both incomplete and incompletely accurate. Only a fraction of functionally important genomic motifs are conserved in syntenic sites; not all are functional; and TFBS often regulate distant genes (Vierstra et al., 2014) (King et al., 2007). Nonetheless, the enrichment of NCCE in genes involved in fundamental developmental processes suggests the NCCE atlas will provide a clear basis for targeted mechanistic analyses and exciting opportunities for future studies.

## ACKNOWLEDGEMENTS

We thank Milladur Rahman, Agata Szade, Sofia Nordling, Mike Lee. Michael Bshneider, Borja Moreno, Yu Zhu, Romain Ballet, Martin Brennan, Jeffrey Ye, Nicole Lazarus, Jean Jang and Dhananjay Wagh for assistance with technical support, cell preparation and single-cell sequencing experiments. We thank Ralf and Susanne Adams for the Ve-Cad5-creERT2 mice and J. Paulson from the The Scripps Research Institute for *St6gal*1^−/−^ mice. This work was supported by NIH grants R01 AI130471 and CA228019, and award I01 BX-002919 from the Department of Veterans Affairs to ECB. TTD was the recipient of a fellowship under NIH training grant T32 AI07290 from the NIAID and T32 HL098049 from the National Heart, Lung, and Blood Institute, and was supported by an AHA Postdoctoral Fellowship and the Stanford Cardiovascular Institute. KB was supported by NIH F32 CA200103. YW was supported by Knut and Alice Wallenberg Foundation KAW 2018.0423. MX was supported by the Tobacco-Related Disease Research Program of University of California, grant T31FT1867. SR and KRH was funded by The New York Stem Cell Foundation (NYSCF-Robertson Investigator to KRH). PB is supported by GINOP 2.3.2-15-2016-00050 and EFOP 3.6.1-16-2016-00004 grants. Graphical abstract and images in Figure 6A were created using BioRender.com.

## AUTHOR CONTRIBUTIONS

TTD conducted or oversaw most experiments. MX performed genomic analyses. AR, JP, YZ, NS, WR, SR, EO, HK, TD and DG performed experiments. PB, FG, BG and GB produced and analyzed the *NKX2-3^-/-^* mice. TTD, AR, JP analyzed experiments. KB generated the ATAC-seq and single-cell RNAseq data. KB and MX analyzed ATAC-seq and single-cell RNAseq data. KRH provided input and advice for experiments. TTD, MX, AR, JP and ECB wrote and edited the manuscript. JP and ECB conceived and supervised the study.

## STAR METHODS

### Lead Contact

Further information and requests for resources and reagents should be directed to and will be fulfilled by the Lead Contact, Eugene C. Butcher.

### Plasmid constructs

mCE-NF**κ**B and mNF**κ**B (−422 and −260 bp upstream of the *Madcam1* transcription start site, respectively) were PCR-amplified as a *Kpn*I-*Hin*dIII fragment from mouse genomic cDNA and cloned into pGL4.11[*luc2P*] to generate the luciferase (LUC) reporter constructs. To analyze the NKX2-3 and NR2F2 binding sites at CE, site-specific mutations were introduced into CE_ΔN-NF**κ**B-LUC via dual PCR and for CE_ΔC_A, and CE_ΔC_AB via Cyagen custom service. CE-LUC, used for control in mutational studies, and human *MADCAM1* reporter constructs hCE-LUC and hCE_NF**κ**B-LUC were made by Cyagen custom service. Control Renilla (Ren) Luciferase vector is from Promega. For co-transfection studies, mammalian expression vectors *Nkx2-3*, *Nr2f2* (COUP-TFII), and *Hey1* were obtained from ABM custom service (Richmond, BC, Canada). All plasmids are described in Supplemental Table S3.

*Nr2f2*, *Hey1*, and *Nkx2-5* cDNA were PCR-amplified as *Bam*HI-*Eco*RI-fragments and *Nkx2-3* and *Nkx2-3*Δ*2-24* cDNA was PCR-amplified as *Eco*RI-*Hin*dIII-fragments from mouse cDNA and cloned into the pET21-A vector (Novagen) to generate His and T7-tagged recombinant proteins. *Nr2f2* cDNA was cloned into the pGEX2TK vector (GE Healthcare) to generate GST-tagged COUP-TFII. pET21A-*Nkx2-1* was from Cyagen, custom service. The fidelity of all constructs was verified by sequencing. Primer sequences are provided in the supplemental material (Table S1).

### ATAC sequencing

Endothelial cells or lymphocytes from BALB/c mice were isolated and subjected to ATAC-seq as previously described (Buenrostro et al., 2015a; Buenrostro et al., 2015b; Lee *et al*., 2014b). Quality control was performed on reads using FastQC (v0.11.9) (http://www.bioinformatics.babraham.ac.uk/projects/fastqc/). Reads were mapped to the mouse reference genome mm10 using Bowtie2 (v2.3.5) (Langmead and Salzberg, 2012) with PCR duplicates removed. Reads were quantified using BEDTools (v2.29.2) (Quinlan and Hall, 2010). Raw read counts were normalized to 1× genome coverage for visualization using the UCSC Genome Browser.

### Cell lines

HEK293T cells were grown in Dulbecco’s modified Eagle’s medium (DMEM) containing high glucose and L-pyruvate and supplemented with 10% heat inactivated fetal bovine serum (FBS) at 37°C and 5% CO_2_. bEnd.3 and DN-MAML bEnd.3 cells were grown in the same media supplemented with penicillin-streptomycin. BEnd.3 and bEnd.3-based transductions were grown and maintaned at confluence until the cells assumed an organized appearance (typically 2-3 days after confluence) prior to stimulation with TNF□.

### Stable transfections

The bEnd.3-DNMAML cell line was constructed by transducing bEnd.3 cells (ATCC CRL-2299) with MSCV-MAML(12-74)-EGFP, which contains amino acids 12-74 of human MAML fused in frame with EGFP (Gift from Warren Pear, University of Pennsylvania). EGFP positive cells were sorted on a BD FACS Aria III. Retrovirus was produced with the Phoenix-Eco packaging cell line (ATCC CRL-3214) using standard methods (Weng et al., 2003).

Stable bEnd.3 DN-MAML cell lines overexpressing *Nkx2-3*, *Nkx2-3* TN domain or control vector, and bEnd.3 cell lines expressing *Hey1*, *Nkx2*-*3*, *Nr2f2*, as well as shRNA for *Nkx2-3*, *Nr2f2* and control shRNA, were generated by transduction with respective retroviral constructs packaged in HEK293T cells via standard protocols (Invitrogen).

### Transient transfection assays

For luciferase reporter assays, 1.5×10^5^ HEK293T cells were plated on 24-well plates followed by transfection after reaching ∼70-80% confluency. The CE_NF**κ**B-LUC and NF**κ**B-LUC constructs were amplified from a pGL4 vector (Promega). CE_ΔN-NF**κ**B-LUC, CE_ΔC_A, CE_ΔC_AB, mouse CE-LUC, and human CE-LUC were amplified from a pGL4.23-based vector custom made by Cyagen (LUC reporter containing a minimal promoter). Renilla (Ren) Luciferase was amplified from a pRL vector (Promega). 36 hours after TNF□ stimulation, cells were harvested by centrifugation and re-suspended in lysis buffer. For Figure 2A, cells were stimulated for 16 hours. Luciferase assays were carried out using the Dual-Glo Luciferase assay system (Promega) and a Turner, TD-20/20 luminometer as previously described (Addepalli and Limbach, 2016). Firefly luciferase values were normalized to Renilla luciferase signal of the TNF stimulated conditions from at least three independent experiments each with 2-3 replicates. Statistical significance was assessed via Student’s one tail or two tail paired t-test.

### Protein expression and purification

The pET21A-*Hey1*, -*Nr2f2*, -*Nkx2-3*, -*Nkx2-3*Δ*2-24*, -*Nkx2-1*, -*NKx2-5* and pGEX2TK-*Nr2f2* protein expression plasmids were transformed into *E. coli* BL21 competent cells. Protein expression and purification were carried out as previously described (Dinh *et al*., 2012) and purified proteins were quantified against BSA.

### Electrophoretic mobility shift assay (EMSA)

EMSAs were performed as previously described (Dinh *et al*., 2012). Probes were generated by annealing 100 pmol of sense and antisense oligonucleotides (Table S1) and 1-2 pmol of probe was used in each reaction. Gel shift reactions were conducted at 4°C in 20% glycerol, 20 mM Tris (pH 8.0), 10 mM KCl, 1 mM DTT, 12.5 ng poly dI/C, 6.25 pmol of random, single-stranded oligonucleotides, BSA and the probe in the amounts specified above. All samples involving only NKX2-3 or HEY1 were loaded on an 8% gel, whereas any involving COUP-TFII were loaded on a 6% gel to resolve protein-DNA complexes. In reactions with cold competitors, 20x unlabeled probes were included in the reactions. Antibodies against specific proteins (anti-NKX2-3, anti-HRT1, Nkx2-1 and Nkx2-5 (Table S2) were from Santa Cruz Biotechnology: sc-83438x, sc-16424x, SC-53136x, and sc-376565X respectively); anti-COUP-TFII from Perseus Proteomics (PP-H7147-00); and anti-T7 antibody from Abcam (ab97964) (Table S2). Antibodies were added to reactions at the same volume amount of the respective protein to obtain super-shifts.

### Chromatin Immunoprecipitation (ChIP)

ChIP was performed using the ChIP-IT High Sensitivity Kit (Active Motif), per manufacturer’s instructions. Cross-linked chromatin isolated from DN-MAML bEnd.3 cells was immunoprecipitated using antibodies against COUP-TFII (Table S2) and the corresponding isotype control. We used PCR primers (Table S1) spanning CE along with a genomic control region (not containing a COUP-TFII binding site). The input and ChIP samples were subjected to real-time PCR (Biorad) and analyzed to ensure that amplification was in the linear range. Three biological samples along with three technical replicates were performed. The data was analyzed as previously described (Dinh et al., 2012). Error bars indicate the standard error of mean. Statistical significance was assessed via two tailed T-test, paired.

### Proximity ligation assay (PLA)

PLA (Sigma) was used to identify specific co-localization of NKX2-3 and COUP-TFII in fixed cells, following manufacturer’s protocols with a few modifications. Prior to the PLA, cells were fixed in acetone at 4°C for 10 min. Cells were incubated with anti-NKX2-3 (Atlas Antibodies, Table S2) and anti-COUP-TFII (Perseus Proteomics; Table S2) or normal rabbit IgG or mouse IgG2a (Santa Cruz Biotechnology) controls. Following PLA, slides were blocked with 10% normal rabbit serum for 30 min at RT and incubated with anti-ERG-488 (Abcam) overnight at 4°C. The slides were then incubated with antibody against Dylight405-conjugated anti-MAdCAM1 (MECA367) for 2 hrs at room temperature and imaged.

### RNA Isolation and Realtime RT-PCR

Total RNA was extracted using TRIzol followed by the RNeasy mini kit (Qiagen) cleanup and RQ1 RNase-free DNase set treatment (Promega) according to the manufacturer’s instructions. First-strand cDNA was synthesized Superscript II (Invitrogen). TaqMan universal master mix reagents (Applied Biosytems) were used for Realtime RT-PCR assays. The primers/probes used in this study are listed in Supplemental Table S1.

### Animals

All experiments were approved by the accredited Department of Laboratory Animal Medicine and the Administrative Panel on Laboratory Animal Care at Stanford and the VA Palo Alto Health Care System (VAPAHCS). COUP-TFII^OE/OE^ (Qin et al., 2013) or COUP-TFII^KO/KO^ (Li *et al*., 2009) mice were crossed with CDH5(PAC)Cre^ERT2^ mice (Sorensen *et al*., 2009) to generate tamoxifen inducible endothelial cell specific COUP-TFII OE or KO mice (EC iCOUP^OE^ ^or^ ^KO^). Littermates not harboring the cre transgene were used as controls. 200µg/g of tamoxifen was injected three times, i.p. every other day.. For the SNA and COUP-TFII^KO/KO^ studies, 100µg/g of tamoxifen was i.p. injected every other day three times. Days post injection correspond to the numbers of days after the first injection. *St6gal1^-/-^* mice (Hennet et al., 1998) were from the Marth lab. *Nkx2-3^-/-^*mice (Pabst *et al*., 1999) were backcrossed to BALB/cJ mice and maintained at the Department of Immunology and Biotechnology, University of Pécs (Czompoly et al., 2011). C57BL/6J mice were obtained from The Jackson Laboratory, Bar Harbor, USA.

### Single cell sequencing

Endothelial cells from the Peyer’s patches of EC-iCOUP^OE^ and control mice were isolated 14-19 days post tamoxifen treatment, dissociated and sorted as previously described (Brulois *et al*., 2020). Single cell gene expression was assayed using the 10x Chromium v3 platform using Chromium Single Cell 3’ Library and Gel Bead Kit v2 (10X Genomics, PN-120237) according to 10X Genomics guidelines. Male and and female cohorts were processed together and resolved post-sequencing. Libraries were sequenced on an Illumnia NextSeq 500 using 150 cycles high output V2 kit (Read 1–26, Read2-98 and Index 1–8 bases). The Cell Ranger package (v3.0.2) was used to align high quality reads to the mm10 transcriptome (quality control reports available: https://stanford.io/37sXZV3). Quality control and data analysis was carried out as described (Brulois *et al*., 2020). Briefly, normalized log expression values were calculated using the scran package (Lun et al., 2016). Imputed expression values were calculated using a customized implementation (https://github.com/kbrulois/magicBatch) of the MAGIC (Markov Affinity-based Graph Imputation of Cells) algorithm (van Dijk et al., 2018) and optimized parameters (t = 2, k = 9, ka = 3). Supervised cell selection was used to remove cells with non-blood endothelial cell gene signatures: lymphatic endothelial cells (Prox1, Lyve1, Pdpn); Pericytes (Itga7, Pdgfrb); fibroblastic reticular cells (Pdpn, Ccl19, Pdgfra); lymphocytes (Ptprc, Cd52). The Arterial (*Gkn3*+), HEC, PCV and CRP clusters were defined based on canonical marker expression. Batch effects from technical replicates were removed using the MNN algorithm (Haghverdi et al., 2018) as implemented in the batchelor package’s (v1.0.1) fastMNN function. Dimensionality reduction was performed using the UMAP algorithm. For UMAP embeddings, cell-cycle effects were removed by splitting the data into dividing and resting cells and using the fastMNN function to align the dividing cells with their resting counterparts. Violin plots were generated using ggplot2; y-axis units for gene expression data correspond to log-transformed normalized counts after imputation. Mean gene expression for male and female cells from each subset were calculated for each sample and plotted with standard errors.

### Identification of conserved NCCE

Conserved genomic regions were defined as regulatory elements with log-odds score greater than 300 in UCSC Genome Browser’s phastCons Placental Elements track, which aligns 60 vertebrate species and measures evolutionary conservation of genomic sequences (Hinrichs et al., 2006; Siepel et al., 2005). COUP-TFII and NKX motifs were searched within each conserved region in the mouse genome using HOMER (Heinz et al., 2010). The spacing between the two motifs within a NCCE was calculated as the distance between the centers of the motifs. Genome coordinates of the motifs were converted from mouse (mm10) to human (hg38) using the liftOver tool (Haeussler et al., 2019), and only mouse coordinates whose syntenic human counterparts correspond to COUP-TFII and NKX binding motifs were defined as conserved NCCE. All NCCEs are provided as a custom track in a UCSC Genome Browser session: https://genome.ucsc.edu/s/m.xiang/NCCE_gap6%2D16_conserved.

### Imaging

PP were imaged following either retro-orbital injection of fluorescent labeled antibodies or immunofluorescence staining of LN sections. Injected antibodies (25–75 µg) were administered 5–30 min prior to sacrifice and PP removal. To image the overall vasculature, the PP was compressed to ∼35–50 µm thickness by applying gentle pressure on a coverslip on a glass slide (Figure 6 and S7-S9). Alternatively, tissues were fixed with 4% paraformaldehyde (PFA), cryoprotected with sucrose, frozen in OCT (Sakura® Finetek) in 2-methylbutane (Sigma) on dry ice and stored at −20 °C. 50 µm cryo-sections were stained with antibodies as previously described (Brulois et al., 2020). Cell lines were fixed with 4% PFA and stained according to standard protocols for imaging.

For *Sambucus nigra* lectin (SNA) studies, binding to the vascular endothelium was assessed by perfusing mice with fluorescently labeled lectin. Briefly, mice were injected iv with a cocktail of fluorescent antibodies to mark endothelial cells and HEV (6μg anti-CD146, 25μg anti-mouse MAdCAM-1 (clone #MECA367), 25μg anti-mouse MAdCAM-1 (clone #MECA89) and 36μg anti-mouse PNAd (clone# MECA79) 15 min prior to anesthesia; followed 5 minutes later by ip injection of 5 units of Heparin (Sigma-Aldrich, H3149). Mice were anesthetized by isoflurane and subjected to vascular perfusion by gravity (75 cm in height) flow with Ca^2+^Mg^2+^ HBSS buffer containing 3 ug/ml SNA (Vector Laboratories). A flow rate of ∼2ml/min was maintained. After perfusion with 20 ml SNA solution, mice were sacrificed, and tissues positioned in Fluoromount-G (Southern Biotech) and manually compressed by firm pressure on an overlying coverslip for confocal imaging. The slides were imaged using Apotome 2.0 fluorescence microscope or LSM 880 laser scanning microscope (Zeiss).

For the *Nkx2-3^-/-^* SNA studies, BALB/c mice were co-housed with *Nkx2-3^-/-^* mice under standard conditions and were used between 6-8 weeks of age. Mice were injected with cocktails containing APC-conjugated anti-CD31 (clone # MEC13.1, BD Biosciences), anti-mouse PNAd (clone #MECA-79) and anti-mouse MAdCAM-1 (clone #MECA-89) labeled with DyLight405 in PBS at 200μl final volume. 20 minutes later the mice were anesthetized with ketamine-xylazine injection ip. supplemented with 0.5 mg/kg atropine. The anesthetized mice were injected with heparin, followed by sequential perfusion through the left ventricle with 10 ml PBS followed by PBS with 2 μg/ml *Sambucus nigra* lectin (SNA) conjugated with Cy3 (Vector Laboratories) containing 0.1mM CaCl_2_ of the period of 20 minutes. After washing with PBS the animals were perfusion-fixed by 4% buffered paraformaldehyde. After fixation, the inguinal lymph nodes and Peyer’s patches were placed onto glass histological slides in an area demarcated with PAP-pen to prevent the spillage of mounting medium. Two strips of double-side sticky tape were placed outside the marked area, and 50 μl of 1:1 mixture of PBS-glycerol was added, in which the samples were immersed, and covered with 22mm x 40mm glass coverslips. The samples were viewed using an Olympus Fluoview FV-1000 laser scanning confocal imaging system (Olympus Europa SE & Co. KG, Hamburg, Germany).

For NKX2-5 and COUP-TFII co-localization studies (Figure S11), BALB/c embryos at E10.0 were isolated in PBS and then fixed in 4% paraformaldehyde (PFA) overnight at 4 °C. The following day, embryos were washed in PBT (PBS containing 0.1% Tween-20), dehydrated in an ascending methanol sequence, xylene treated, and embedded in paraffin. Immunofluorescence was performed on 7 µm deparaffinized sections. Briefly, sections were subjected to antigen retrieval in Tris buffer pH 10.2 for 8 min, washed in 0.1% PBT and incubated in blocking buffer (0.2% milk powder, 99.8% PBT) for 1 h at room temperature. Primary antibodies were incubated in blocking buffer overnight at 4 °C. The following day, the sections were washed three times with PBT and incubated for 2 h with corresponding secondary antibodies in blocking buffer at room temperature. After three washes in PBS, DAPI (Sigma-Aldrich) was added to counterstain the nuclei. The sections were mounted using Prolong Gold Antifade Reagent (Invitrogen) and imaged using Zeiss LSM-700 confocal microscope. The following primary antibodies were used: ERG (Abcam), NKX2-5 (Santa Cruz), and COUP-TFII (Perseus Proteomics). Secondary antibodies were Alexa Fluor conjugates 488, 555, and 647 (Life Technologies).

## SUPPLEMENTAL TABLES

**Table S1. Primers and probes**

**Table S2. List of Antibodies**

**Table S3. List of plasmids**

**Table S4. NCCE genomic loci**

---

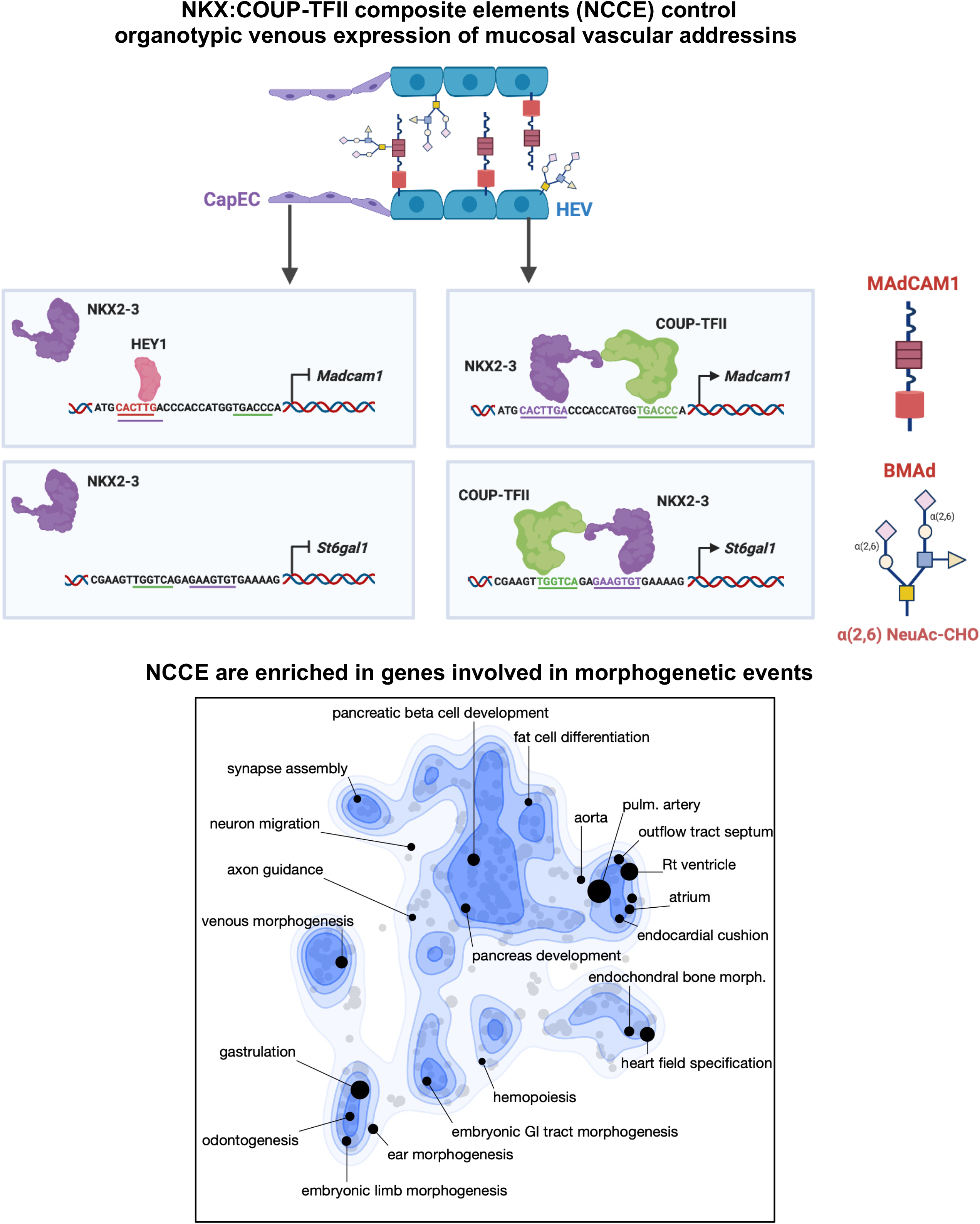

**Figure S1.**
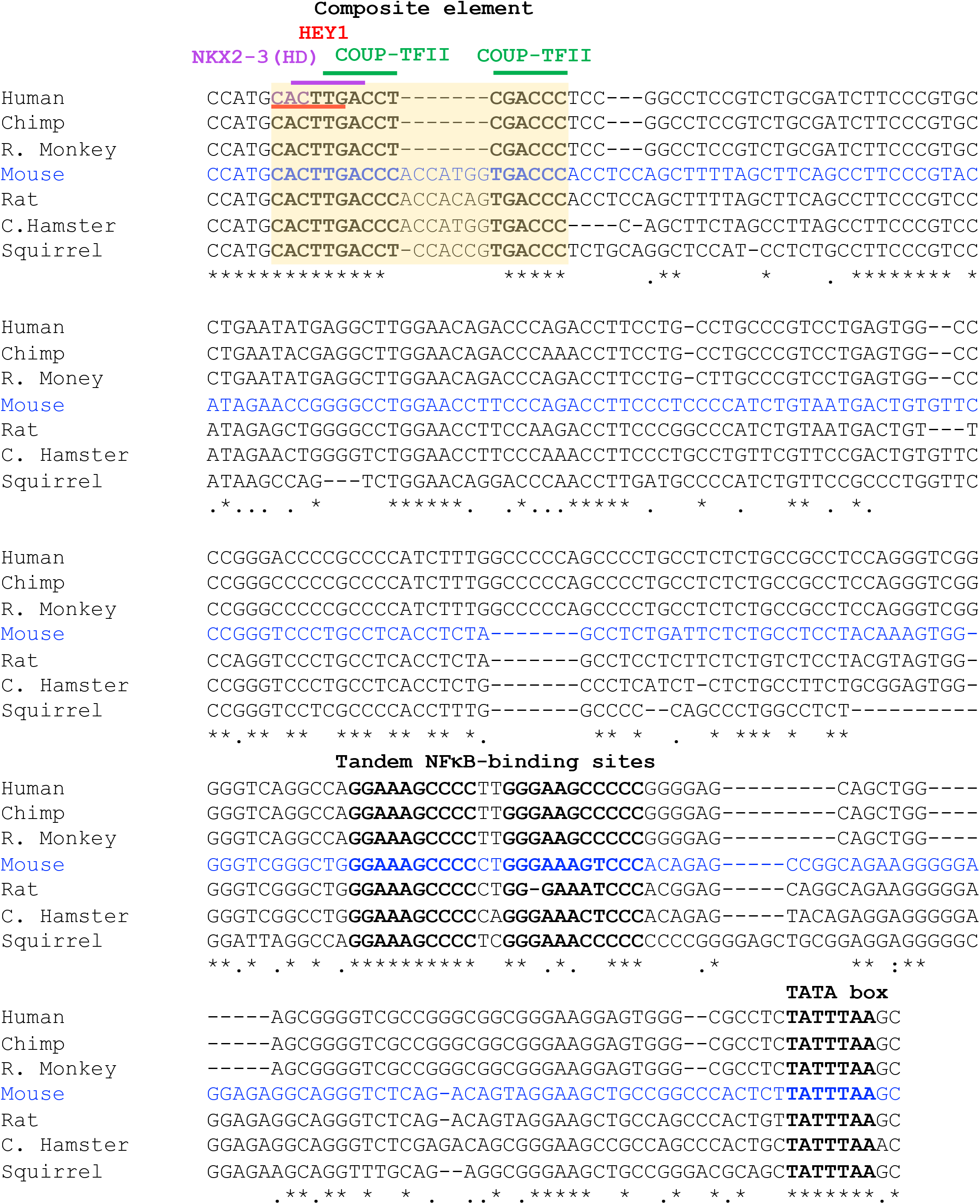
Conserved transcription factor binding sites in the *Madcam1* promoter region. Conserved binding sites for transcription factors NKX2-3 (purple line; HD), HEY1 (red underline) and COUP-TFII (green line) within the NKX2-3/COUP-TFII composite element (CE; yellow box), and proximal NF**κ**B sites are shown. Conserved nucleotides are marked by an asterisk (*). The sequences included in mouse CE-NF**κ**B LUC reporter constructs are shown in blue.

**Figure S2.**
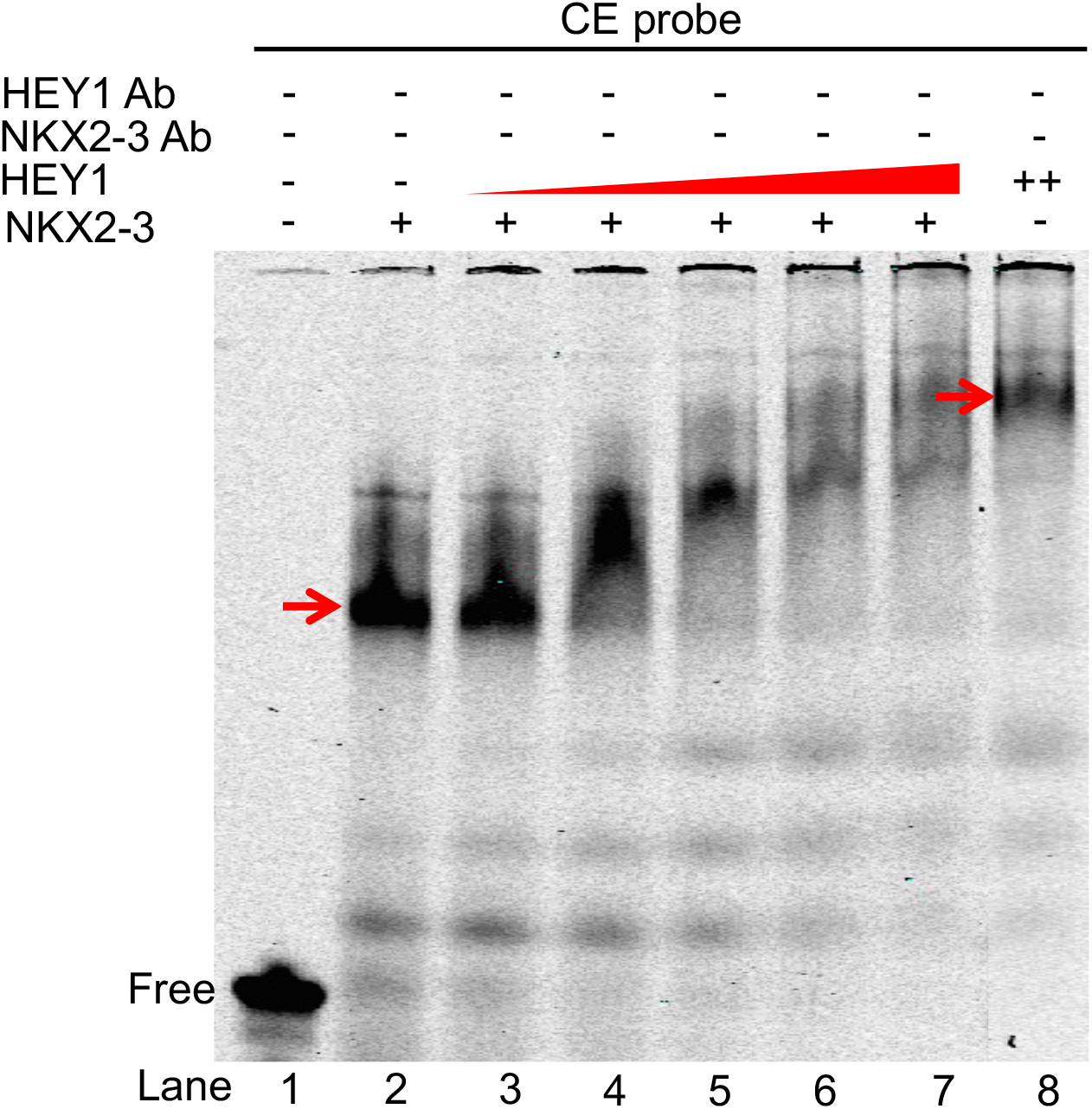
HEY1 competes with NKX2-3 for binding to CE. EMSA showing migration of recombinant HEY1 and NKX2-3 bound to the CE probe (arrows). Progressive shift in migration indicates displacement of NKX2-3 by co-incubation with increasing concentrations of HEY1.

**Figure S3.**
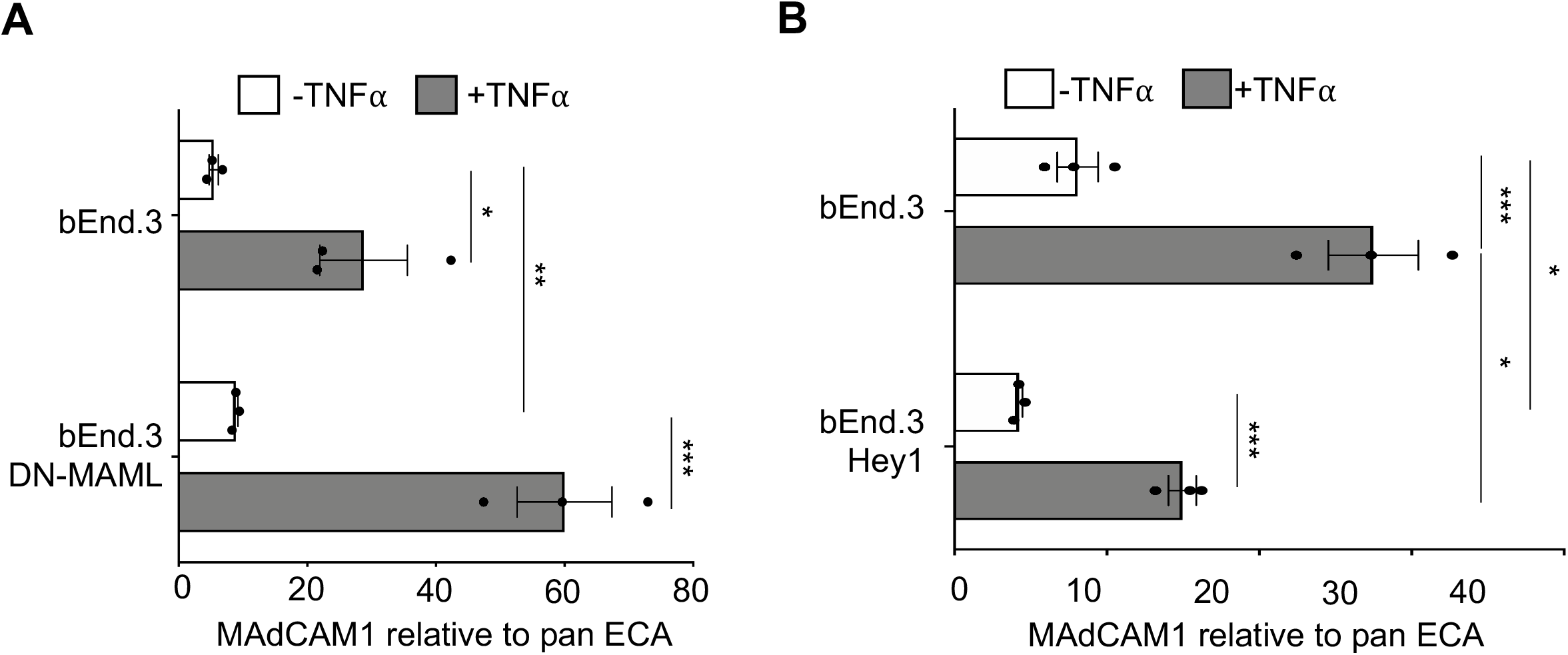
Notch signaling inhibits endothelial cell surface MAdCAM1 expression. ELISA of MAdCAM1 in bEnd.3 vs (A) DNMAML-bENd.3 cells or (B) Hey1-bEnd.3 cells. Control unstimulated and 24 hr TNF□-stimulated cells are denoted by white and gray bars, respectively. Data are normalized to binding of anti-PLVAP antibody MECA32, a pan-EC marker, and shown as mean of three biological replicates with SEM. ***: p-value <0.005, **: p-value <0.01, *: p-value<0.05 by a two tailed t-test.

**Figure S4.**
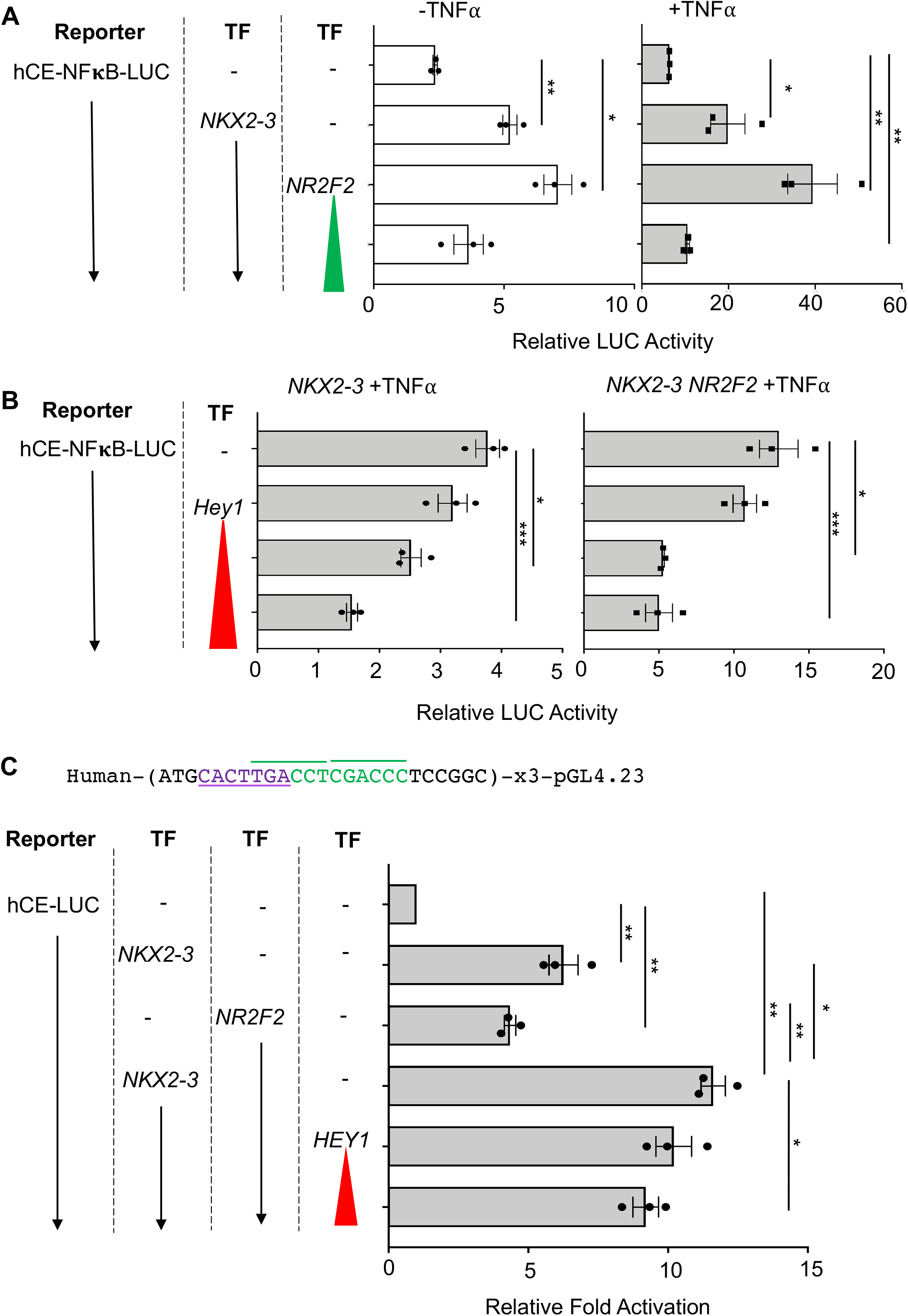
Functional properties of *MADCAM1* CE in human. (A) Activity of luciferase reporter driven by CE-containing human *MADCAM1* promoter is enhanced by *NKX2-3* cooperatively with *NR2F2* in 293T cells. (B) HEY1 dose-dependently suppresses activation of the *MADCAM1* reporter, when co-transfected with *NKX2-3* (left) or with *NKX2-3* and *NR2F2* (right) in 293T cells. (C) Activity of luciferase reporter driven by promoter with three copies of the human *MADCAM1* CE sequence fused in tandem, when co-expressed with *NKX2-3*, *NR2F2* and *HEY1* in 293T cells. All data are mean ± SEM from three independent experiments, each with 3 technical replicates. *: p-value<0.05, **: p-value <0.01, ***: p-value <0.005; two tailed t-tests, paired.

**Figure S5.**
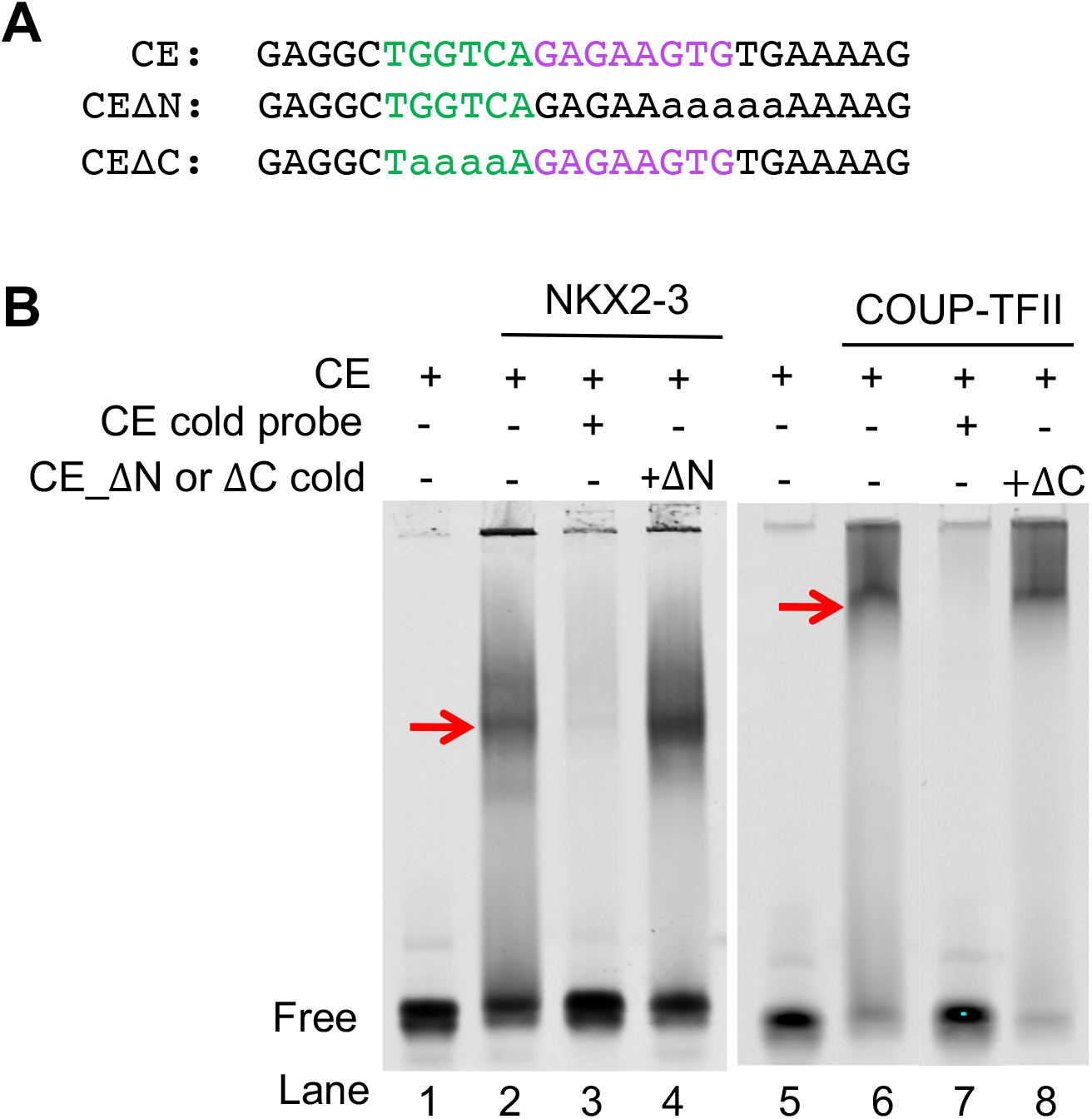
Role of NKX2-3 and a conserved NCCE in regulation of *St6gal1*. (A) Sequence of wildtype and mutant CE probes for EMSA. CE_ΔN harbors mutations in the NKX2-3 binding site; CE_ΔC harbors mutations in the COUP-TFII binding site. (B) EMSA illustrating NKX2-3-HIS and COUP-TFII-GST recombinant proteins binding to the St6gal1 WT probe (red arrows). Cold WT probe outcompeted the CE probe, while mutated probes (CE_ΔN or CE_ΔC) failed to compete against the WT probe.

**Figure S6.**
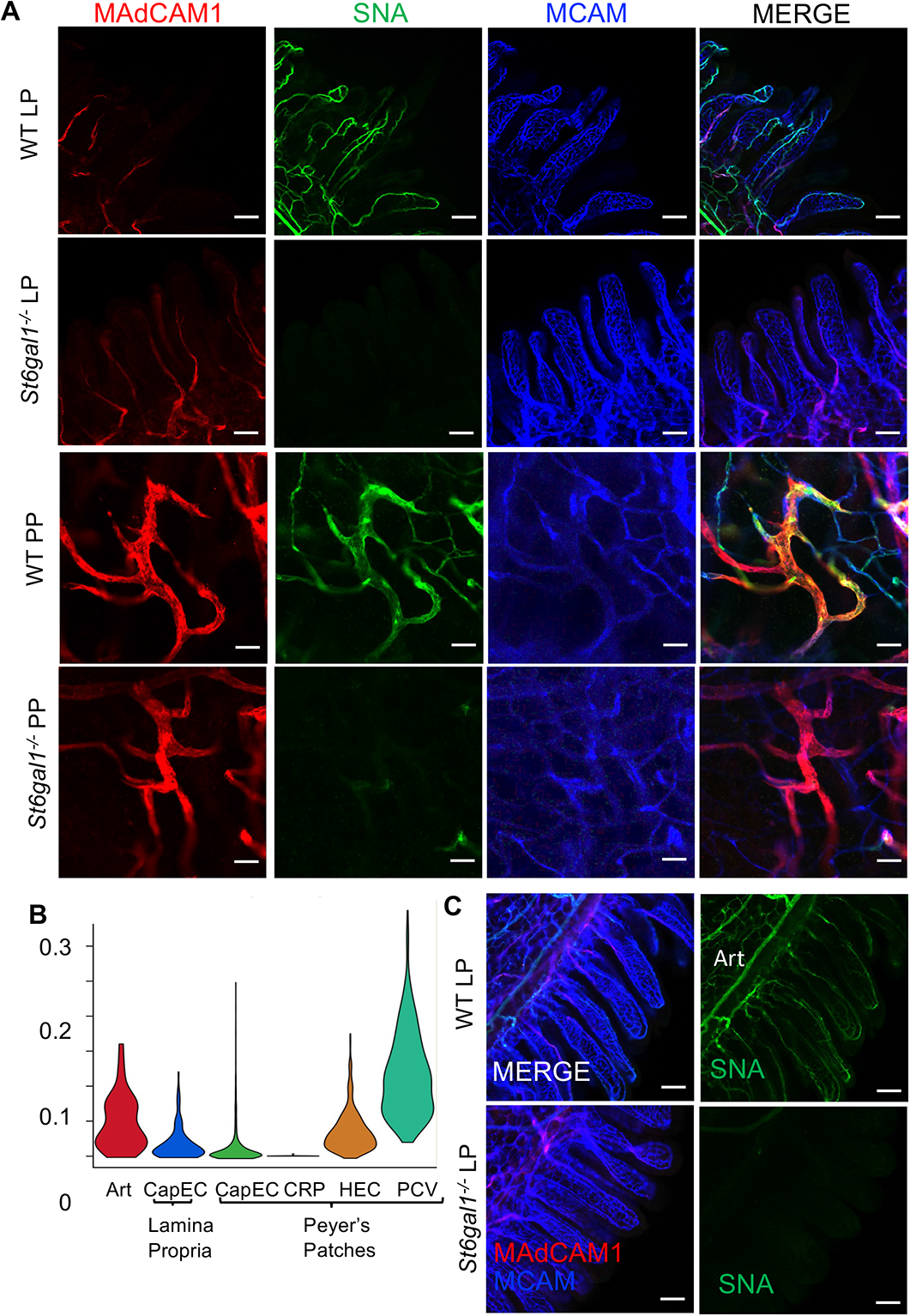
*St6gal-*dependent SNA binding to vascular endothelium in PP and LP. (A) Binding of the alpha-2,6-sialic acid-specific lectin SNA to the LP and PP HEV in WT vs *St6gal1^-/-^* mice. Mice were injected with pan-EC anti-MCAM (CD146, blue) and AF450-labeled anti-addressin antibodies MECA89 and MECA367 to label HEV (red), followed by vascular prefusion with AF488-labeled SNA lectin (green). Scale bars: 100µm, 10x (top two rows), 50µm, 20x (bottom two rows); whole mount imaging. (B) *St6gal* expression in select EC subsets in LP and PP, assessed by scRNAseq. (C) SNA stains the artery (Art) in WT but not *St6gal1^-/-^* mice. Whole mount, 10x, scale bars: 100µm.

**Figure S7.**
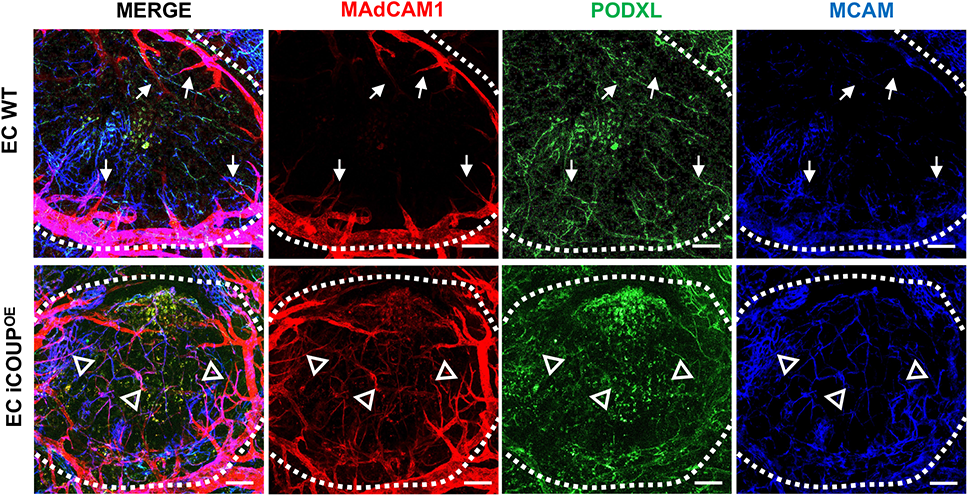
MAdCAM1 expression extends into the PP follicle in EC-iCOUP^OE^ mice. Confocal image showing MAdCAM1 extension from HEV into capillaries within the PP of EC-iCOUP^OE^ (arrowheads) but not control WT mice (arrows). Scale bars: 100µm, 20x, whole mount, 14 dpi.

**Figure S8.**
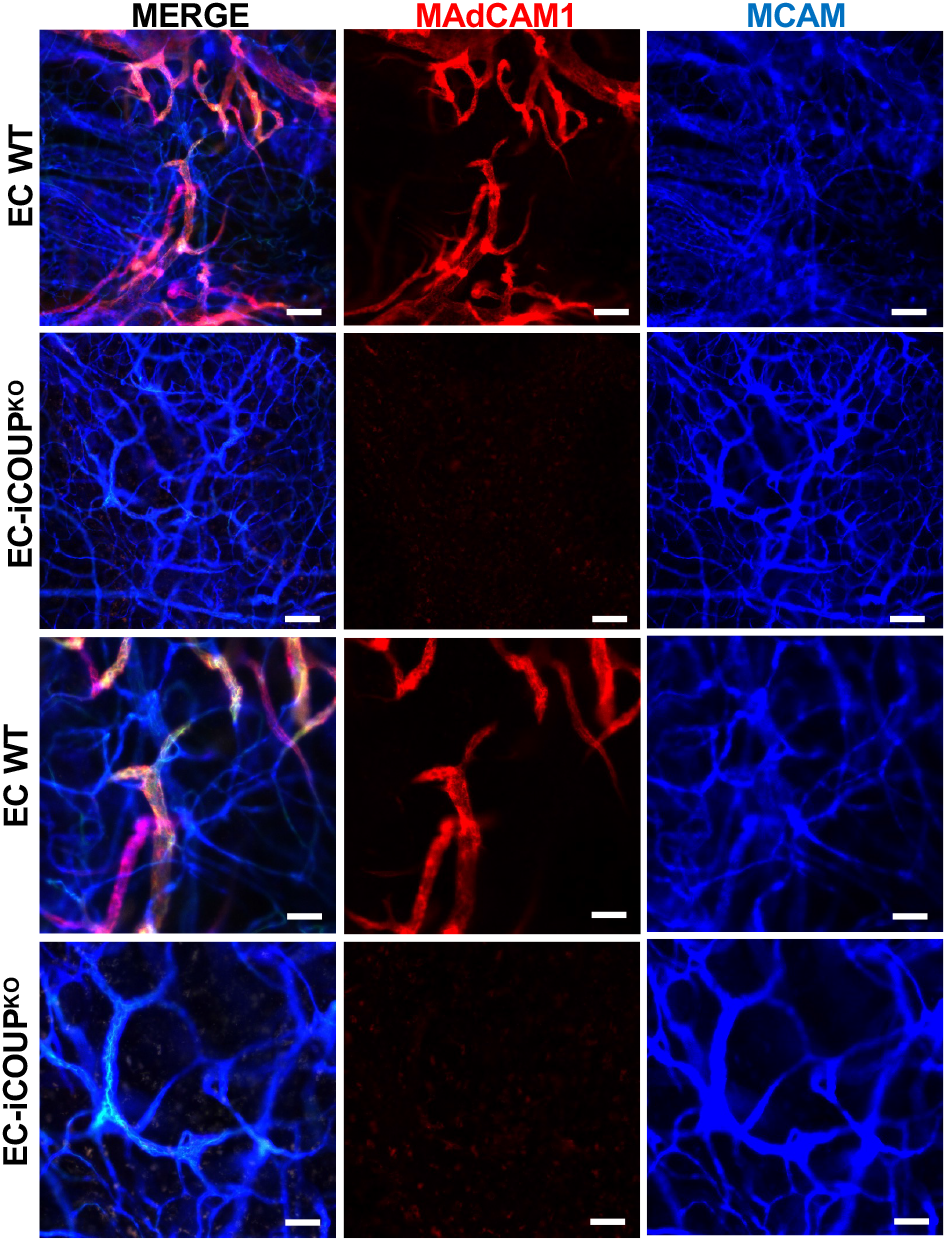
Loss of MAdCAM1 in the PP HEC of iCOUP^KO^ mice. Confocal image showing distinct patterns of loss of MAdCAM1 in PP HEC in EC-iCOUP^KO^ vs control WT mice. EC stained by i.v. injection of directly conjugated antibodies as in Fig S6. Scale bars: 100µm, 10x (top two rows), 50µm, 20x (bottom two rows); whole mount imaging.

**Figure S9.**
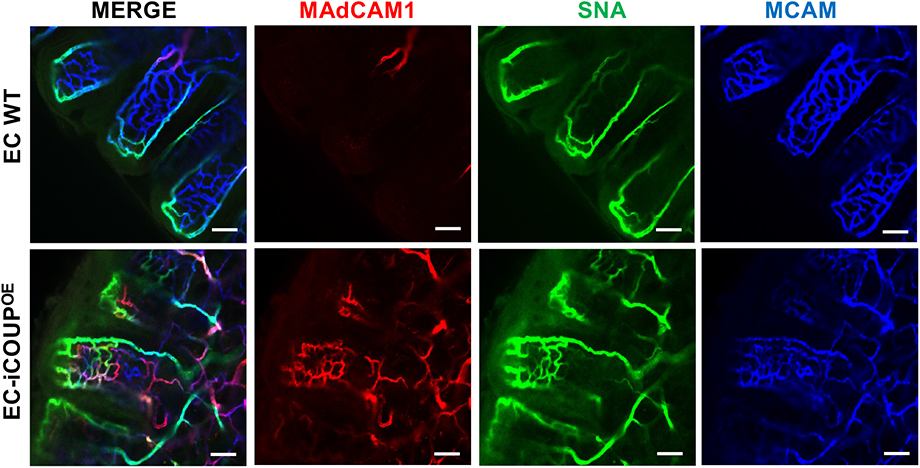
Ectopic co-expression of MAdCAM1 and SNA-binding glycotopes in the villus capillary network in EC-iCOUP^OE^ mice. Confocal image showing distinct patterns of MAdCAM1 staining and SNA binding by lamina propria capillaries in EC-iCOUP^OE^ vs control WT mice. Scale bars: 50µm, 20x, whole mount, 19 dpi.

**Figure S10.**
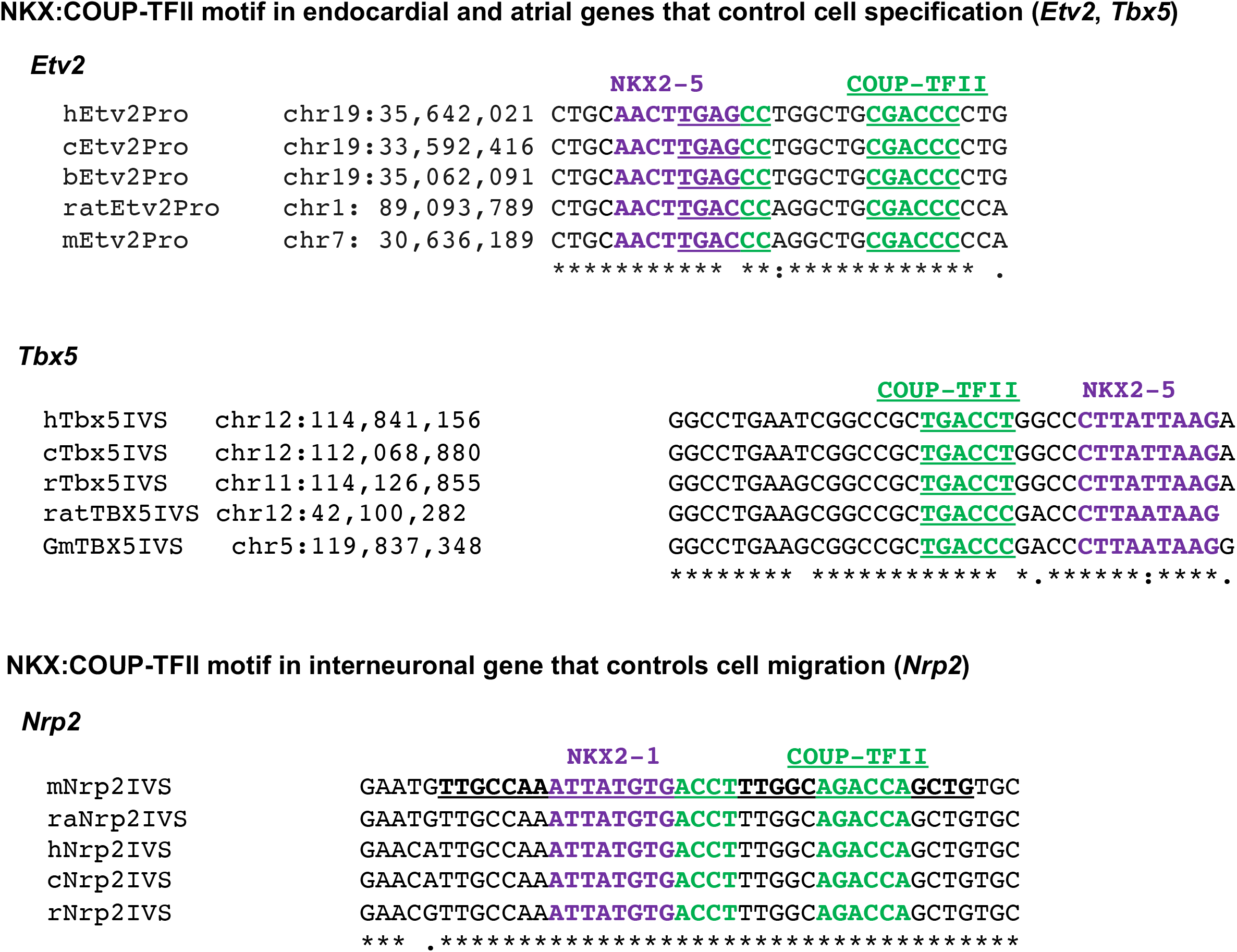
Conserved NCCE in select genes controlled by NKX and COUP-TFII in cardiac or neuronal development. Alignment of candidate NCCE sequences from the promoter region of *Etv2* or the intronic regions of *Tbx5* and *Nrp2* across species. h, human; c, chimpanzee; b, baboon; r, rhesus monkey; ra, rat; m, mouse. Pro, promoter; IVS, intron. COUP-TFII and NKX family HD binding sites are highlighted in green and purple, respectively.

**Figure S11.**
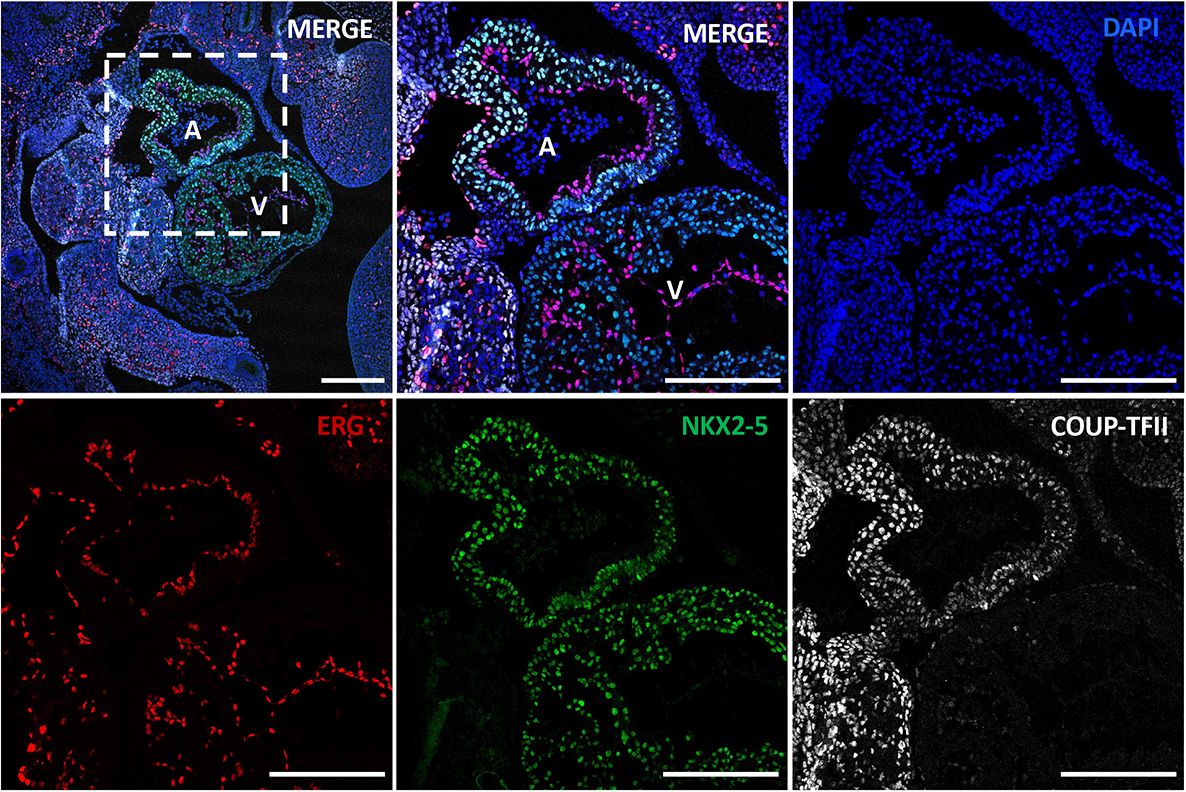
NKX2-5 and COUP-TFII co-localize in the developing mouse atrium. Representative image of mouse heart (e10) stained for DAPI (blue), ERG (red), NKX2-5 (green) and COUP-TFII (white). Scale bars: 200 µm. A, atrium; V, ventricle.

**Figure S12.**
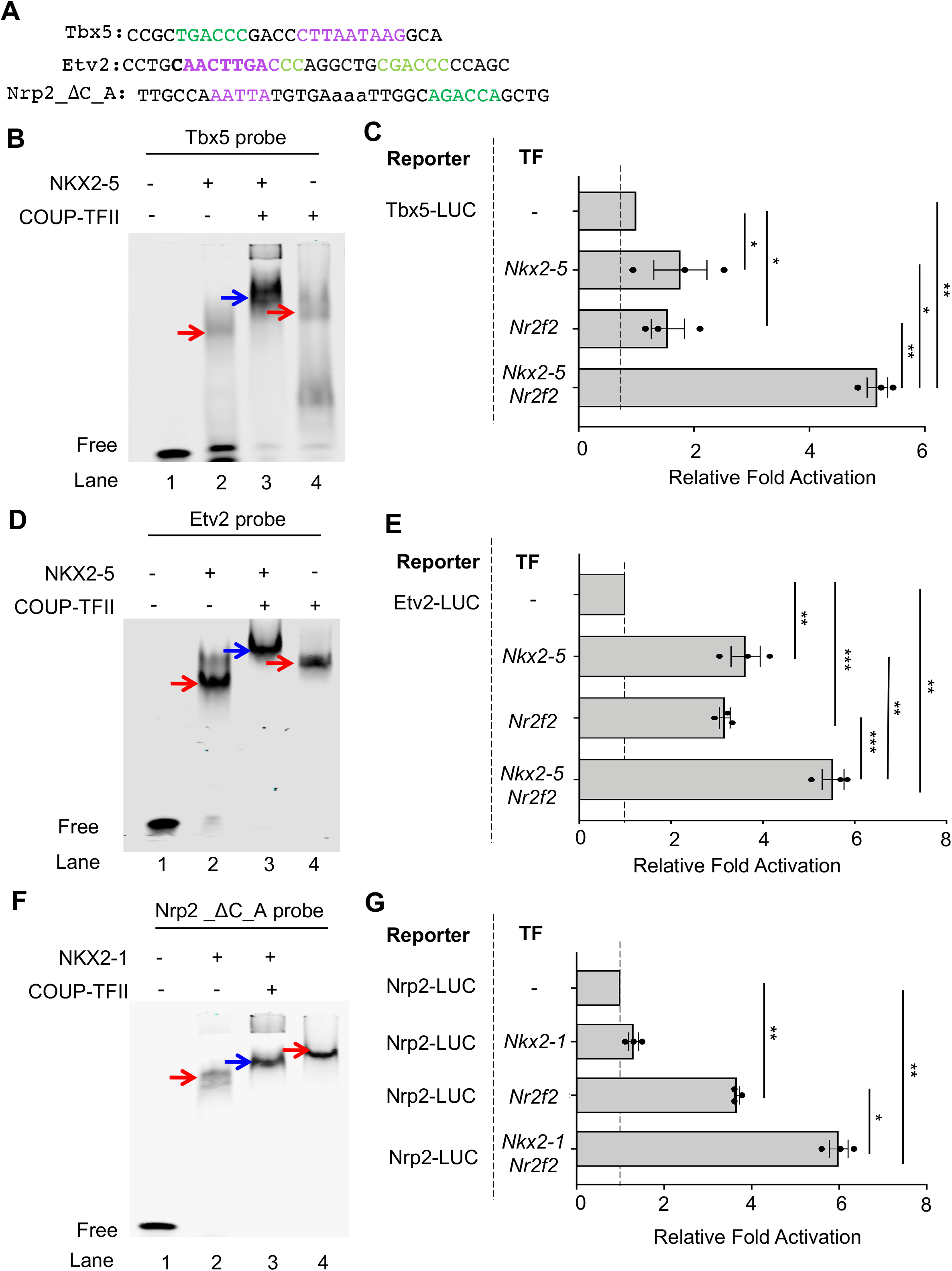
Conserved NCCE in *Tbx5* and *Etv2* promoters support combinatorial control of transcription by NKX2-5 or NKX2-1 and COUP-TFII. (A) Sequence of probes used for EMSA. (B, D, F) Binding of recombinant COUP-TFII and NKX2-5 or NKX2-1 to the *Tbx5* (B), *Etv2* (D) and *Nrp2* (F) NCCE probes. Heterodimeric complexes are indicated by blue arrows. (C, E, G) Activity of luciferase reporters driven by NCCE-containing Tbx5 (C), Etv2 (E) or Nrp2 (G) promoter, with or without *Nr2f2*, *Nkx2-5* or *Nkx2-1*. All data are mean ± SEM from three independent experiments, each with 3 technical replicates. *: p-value<0.05, **: p-value <0.01, ***: p-value <0.005; two tailed t-tests, paired.

